# Synthetic gene circuits that selectively target RAS-driven cancers

**DOI:** 10.1101/2024.11.11.622942

**Authors:** Gabriel Senn, Leon Nissen, Yaakov Benenson

## Abstract

Therapies targeting mutated RAS, the most frequently mutated oncogene in human cancers, could benefit millions of patients. Recently approved RAS inhibitors represent a breakthrough, but are limited to a specific KRAS^G12C^ mutation and prone to resistance. Synthetic gene circuits offer a promising alternative by sensing and integrating cancer-specific biomolecular inputs, including mutated RAS, to selectively express therapeutic proteins in cancer cells. A key challenge for these circuits is achieving high cancer selectivity to prevent toxicity in healthy cells. To address this challenge, we present a novel approach combining multiple RAS sensors into RAS-targeting gene circuits, which allowed us to express an output protein in cells with mutated RAS with unprecedented selectivity. We implemented a modular design strategy and modelled the impact of individual circuit components on output expression. This enabled cell-line specific adaptation of the circuits to optimize selectivity and fine-tune expression. We further demonstrate the targeting capabilities of the circuits by employing them in different RAS-driven cancer cells and provide evidence for their therapeutic potential by linking them to the expression of a clinically relevant output protein, which induced robust killing of cancer cells with mutated RAS. This work highlights the potential of synthetic gene circuits as a novel therapeutic strategy for RAS-driven cancers, advancing the application of synthetic biology in oncology.

## Introduction

RAS (Rat sacroma) mutations are the most common oncogenic alterations in human cancers^1^, accounting for 19% of all cases^2^. The RAS gene family consists of HRAS, KRAS and NRAS, with KRAS being the most frequently mutated isoform^3^. Considered “undruggable” for many years, recent approval of several KRAS^G12C^ inhibitors^4^ has demonstrated the efficacy of targeting RAS in cancer treatment. However, these therapies are limited to cancers with KRAS^G12C^ mutations, which narrows the target range and makes the inhibitors susceptible to resistance development due to the emergence of other RAS mutations^5^. While KRAS^G12C^ remains the only RAS mutation with approved targeted inhibitors, recent advances have expanded the therapeutic landscape. Several inhibitors targeting other RAS mutations, including KRAS^G12D^ (e.g., MRTX1133)^6^, as well as pan-RAS inhibitors^7^, are currently in clinical development. A comprehensive overview of these efforts is provided by Oya and colleagues 2024^8^.

Additional RAS-targeting strategies have encompassed protein engineering, RAS silencing, and artificial promoters. Synthetic proteins and endogenous RAS effectors (proteins that interact with active RAS and transduce its signaling) were engineered to bind^9^, inhibit^10^, or degrade RAS^11,12^. Additionally, miRNAs modulated by oncogenic RAS were identified as potential drug targets^13^ and siRNA against KRAS was used to target RAS-driven cancers in mice^14^. Artificial promoters or synthetic transcription factor response elements responsive for transcription factors downstream of RAS were developed to interrogate RAS signaling^15^ and target cancer cells with RAS mutations by expressing a toxin-antitoxin system ^16,17^.

Gene circuits that logically integrate multiple biomolecular inputs are an emergent modality that allows discriminating between malignant and healthy cells and selectively eliciting a therapeutic response only in target cells^18–20^. Logic gene circuit-based therapeutic prototypes already demonstrated efficacy and safety in animal tumor models^21^. Some of the previously-explored approaches for sensing and targeting RAS^13,15–17,22^ are potentially compatible with the logic gene circuit paradigm. Two circuits have already been shown to build upon, or directly sense, RAS signaling. Gao and colleagues demonstrated a split protease approach to sense over-activation of proteins upstream of RAS. By fusing one domain of a protease to HRAS and the other domain to a RAS binding domain (RBD) of rapidly accelerated fibrosarcoma type C (CRAF), they created a sensor for overexpressed mutated epidermal growth factor receptor (EGFR) or son of sevenless 1 (Sos-1) in HEK293 cells^23^. Vlahos and colleagues adapted this approach to directly sense RAS activation by fusing either part of a split protease to an RBD domain, enabling RAS-dependent interleukin-12 secretion in HEK293T cells overexpressing KRASG12V.^24^ Further advancing synthetic circuits for therapeutic applications against RAS-driven cancer requires achieving high selectivity for mutated RAS and robust function in heterogeneous cancer cells. High selectivity is required to achieve strong output expression in cancer cells while minimizing off-target effects and toxicity in healthy cells with wild-type RAS. In addition, eventual translatability of these approaches to the clinic requires improved robustness in the face of tumor heterogeneity.

In this study, we address these challenges by developing a set of versatile RAS sensors and RAS-dependent transcription factor response elements to create synthetic gene circuits that use RAS activation as input. By combining these direct and indirect RAS sensors in an AND-gate configuration, we develop RAS-targeting circuits with high dynamic range and high selectivity towards cells with mutant RAS. The modular design allows fine-tuning the circuits to specific target and off-target cells and balancing their activation strength versus leakiness. Finally, we demonstrate that our new RAS-targeting circuits function as cancer cell classifiers, selectively expressing an output protein in a wide range of cancer cell lines with RAS-overactivating mutations while maintaining minimal output in cell lines with wild-type RAS.

## Results

### Design of a synthetic RAS sensor

The endogenous RAS signaling pathway is initiated by the binding of cytoplasmic effector proteins to activated RAS-GTP. The effectors sense RAS-GTP dimerization and propagate the signal downstream through a phosphorylation cascade. Post-activation, GTPase-activating proteins, such as neurofibromin 1 (NF-1), facilitate the hydrolysis of RAS-bound GTP to GDP, thereby terminating the signaling. Cancer-associated mutations in RAS render it insensitive to this hydrolysis, resulting in constitutively active RAS^25^ that drives uncontrolled cell proliferation and tumor growth.

Inspired by the natural function of CRAF, a key effector of RAS^26^, we designed a sensor that exploits the selective binding to RAS-GTP of CRAF’s RBDCRD (**R**AS-**b**inding **d**omain/**c**ysteine **r**ich **d**omain) domain. In our sensor, we fuse the RBDCRD domain to engineered truncated and mutated NarX variants originally derived from the bacterial two-component system, namely NarX^379–598^H399Q and NarX^379-598^N509A (Fig. 1a). We showed previously that these NarX variants are able to transphosphorylate in mammalian cells, but only upon forced dimerization via fused protein domains. Therefore, NarX variants can functionally replace CRAF’s own dimerization domain when fused to RBDCRD, while enabling orthogonal signaling in mammalian cells via a humanized NarL response regulator^27^. We call these chimeric constructs “RBDCRD-NarX fusions”.

**Figure 1.**
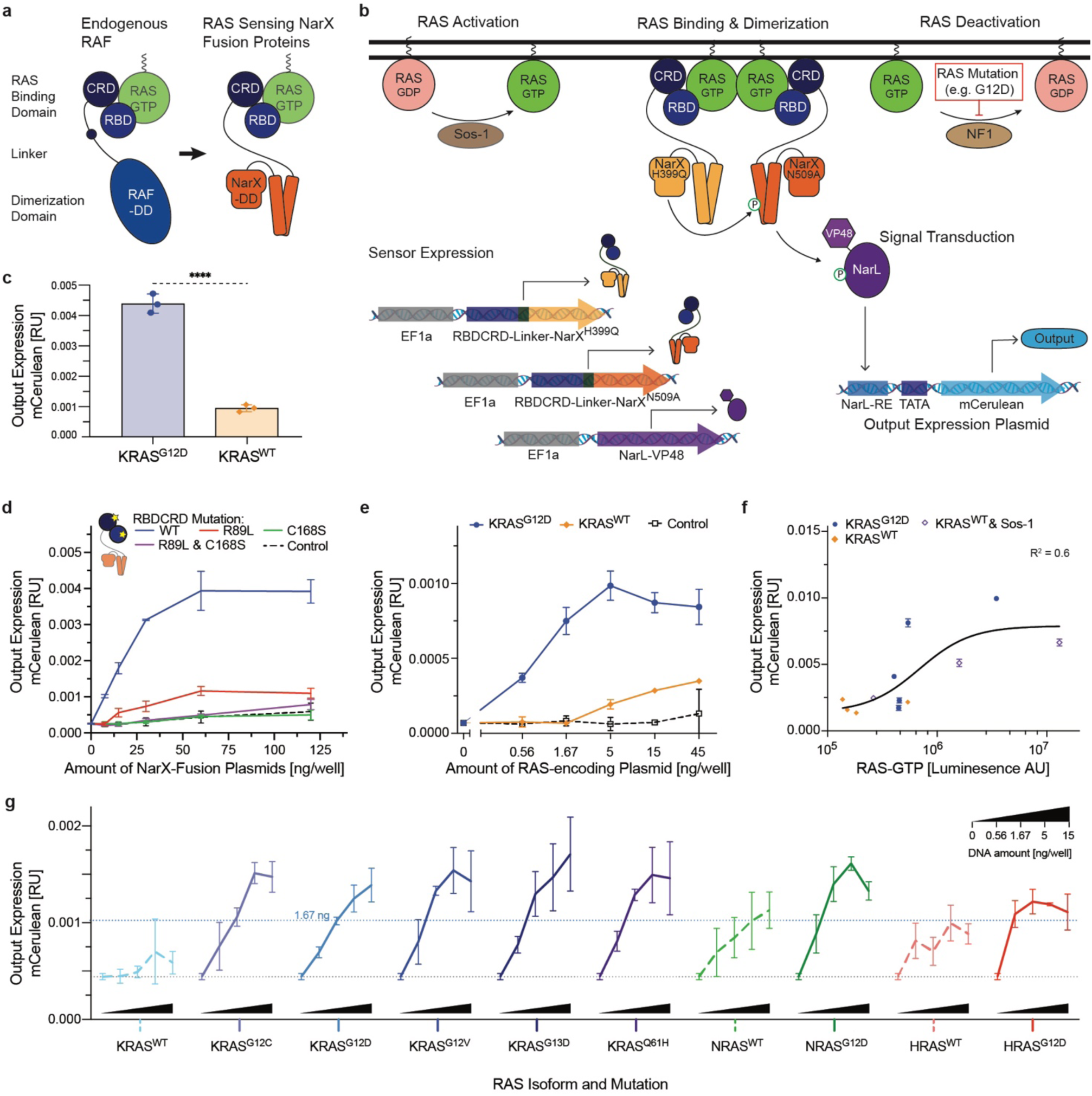
Design and Characterization of the RAS Sensor. (**a**) Design of the binding component of the RAS sensor. Inspired by natural RAF (left), the RAS-binding component of the sensor (right) comprises RAS binding domain (RBDCRD), a linker, and a NarX-derived transphosphorylation domain. (**b**) Schematic of the RAS sensor composition and mechanism of action. The sensor’s genetic payload is encoded on four plasmids. Two plasmids express the RAS-binding components: RBDCRD fused, respectively, to NarX^N509A^ or NarX^H399Q^. The third plasmid expresses NarL-VP48, and the fourth plasmid encodes the output protein (mCerulean) under the control of a NarL response element (NarL-RE) in front of a minimal promoter (TATA). Upon RAS activation, the RBDCRD domain of the RAS-binding components bind to RAS-GTP. This binding leads to a forced dimerization of the NarX domains and a transphosphorylation of NarX^N509A^, in turn phorsphorylating NarL. Phosphorylated NarL binds its response element on the output plasmid, inducing the expression of the output protein. (**c**) Sensor activation by mutated RAS. The bar chart shows output expression in HEK293 cells co-transfected with the RAS Sensor and either KRAS^G12D^ or KRAS^WT^. (**d**) Dose-response curve and dependence of the RAS sensor on functional RAS binding. Output expression of RAS sensors with either RBDCRD wild-type (blue) or RBDCRD with R89L (red), C168S (green), or both (purple) mutations. The dashed line represents conditions where the NarX-fusion plasmids were replaced with a non-coding plasmid (control). (**e**) Dependence of sensor output on RAS levels. Output expression of the RAS sensor measured with increasing amounts of KRAS^G12D^ (blue), KRAS^WT^ (orange), or negative control (black) plasmid. (**f**) Input-output curve. Correlation of the output expression with the RAS-GTP levels in HEK293 cells, measured by a luminescence RAS-pulldown ELISA assay. To alter RAS-GTP levels, the cells were transfected with different amounts of either KRAS^G12D^ (blue), KRAS^WT^ (orange), or KRAS^WT^ + Sos-1 (purple) plasmids. Pearson’s correlation is shown as R^2^. **(g)** Generalizability across RAS variants. Output expression of the RAS sensor when co-transfecting increasing amounts of different RAS isoforms and mutants. mCerulean output expression was measured by flow cytometry and normalized to a constitutively expressed mCherry transfection control. Mean values were calculated from biological triplicates. Error bars represent +/- SD. Significance was tested using an unpaired two-tailed Student’s t-test. ****p < 0.0001.

We surmised that the RAS sensor would function as follows: activation of RAS (whether endogenous or mutation-driven) would elevate RAS-GTP levels, which would in turn bind the RBDCRD domain of the RBDCRD-NarX fusion. Resulting dimerization of RBDCRD would lead to a forced dimerization of the fused NarX, leading to NarX^H399Q^ transphosphorylating NarX^N509A^, in turn phosphorylating NarL. Phosphorylated NarL would bind to its response element on the NarL-responsive promoter and induce the expression of an output protein, here mCerulean (Fig. 1b). In order to test these assumptions, we constructed the necessary components including the two complementary RBDCRD-NarX fusions, the humanized NarL and the NarL-reponsive promoter coupled to mCerulean. Initially, we chose HEK293 cells as a test system. To emulate the presence of mutant RAS we transfected HEK293 cells with a plasmid expressing KRAS^G12D^, and to emulate high levels of wild-type RAS we transfected them with a plasmid-encoded wild-type KRAS (KRAS^WT^). (Note that HEK293 cells express low endogenous levels of wild-type KRAS, HRAS and NRAS.) In the first set of tests, delivery of all the sensor components to HEK293 cells using transient transfection showed significantly higher output expression in cells expressing KRAS^G12D^ than in cells transfected with KRAS^WT^ (Fig. 1c). In HEK293 cells transformed with the mutant KRAS, sensor response increased upon increase in the dose of the sensor-encoding plasmids (Fig. 1d). Further, sensor function depended on RAS binding, because mutations in the RAS binding domain (RBD) and the cysteine-rich domain (CRD) strongly decreased output expression. To demonstrate this, we mutated two residues in RBDCRD important for RAS-RAF signaling: Arginine 89 in RBD^28^ and Cystein 168 in CRD^29^. Arginine 89 forms electrostatic interactions with acidic residues in the RAS switch I region, stabilizing the RBD–RAS complex. Cystein 168 is part of a zinc finger motif that is critical for high-affinity association with RAS and efficient RAF signaling ^30,31^. Mutating these residues – R89L in RBD or C168S in CRD– has been shown to reduce RAS binding, diminish RAF kinase activity^30,31^, and impair RAS-dependent membrane localization of RBD–CRD fusion proteins^32^. Interestingly, while the RBD^R89L^ mutant still exhibited dose-dependent leakiness, the CRD^C168S^ and the double RBD^R89L^CRD^C168S^ mutant reduced output expression to background levels (Fig. 1d). In agreement with previous reports^30^ and with our observation that the sensor employing RBDCRD instead of RBD alone shows increased downstream signaling (Supplementary Fig. 1), this result confirms that both RBD and CRD domains are involved in RAS binding and sensor activation.

We then characterized sensor response to increasing input levels. As expected, increased levels of mutant KRAS drove higher output expression, whereas wild-type KRAS resulted in some sensor activation but at much higher expression levels (Fig. 1e). Finally, we measured the RAS-GTP protein levels in HEK293 cells using a RAS-pulldown ELISA assay. We manipulated the RAS-GTP levels in the cells by co-expressing different amounts of KRAS^WT^, KRAS^G12D^ or KRAS^WT^ + Sos-1, a guanine nucleotide exchange factor that activates RAS, and were able to directly correlate higher RAS-GTP levels with higher output expression (Fig. 1f).

To assess the generalizability of the RAS sensor across different oncogenic mutations and RAS isoforms, we tested a panel of oncogenic RAS variants, including multiple KRAS mutants as well as HRAS^G12D^ and NRAS^G12D^. In all cases, we observed concentration-dependent activation of the sensor (Fig. 1g). Among the KRAS mutants, no significant differences in sensor output were observed (Supplementary Fig. 2a–b), suggesting that the behavior characterized with KRAS^G12D^ is representative of other activating KRAS mutations. In contrast, we found more heterogeneous responses across RAS isoforms (Supplementary Fig. 2c–f). Notably, high overexpression of wild-type HRAS or NRAS resulted in stronger sensor activation than wild-type KRAS (Supplementary Fig. 2f). This indicates that all wild-type RAS isoforms can activate the sensor, underscoring the need to consider physiological activity of all isoforms when evaluating circuit specificity and minimizing potential off-target effects in healthy cells.

Collectively, this dataset provides evidence that the RAS sensor is activated by RAS-GTP, and therefore can selectively sense mutant RAS variants because the hallmark of these mutants are increased RAS-GTP levels. Nonetheless, wild-type RAS also binds GTP, which explains sensor response in HEK293 cells that overexpress wild-type RAS and contain relatively high levels of RAS-GTP (Fig. 1f).

### Mechanism of Action

To recapitulate, our design anticipates that the function of the RAS sensor requires the following steps: delivery and expression of the RAS sensor components; RAS-GTP binding of RBDCRD-NarX fusion proteins via their RBDCRD domain, resulting in forced dimerization of the NarX domains and transphosphorylation; NarL phosphorylation; and NarL-mediated output expression. To quantify the effect of RAS activation on these steps, we manipulated the RAS-GTP levels in HEK293 cells in a variety of ways: (i) we overexpressed wild-type KRAS to show the effect of high non-mutant KRAS concentration on downstream sensor response and quantify sensor activation in the presence of RAS-GTP associated with wild-type KRAS; (ii) we overexpressed mutant KRAS in order to measure sensor response to the mutant KRAS input; (iii) we overexpressed NF1, an endogenous GTPase-activating protein that deactivates residual endogenous wild-type RAS in HEK293 cells and allows to quantify non-specific sensor response in the absence of RAS-GTP; and (iv) we expressed Sos-1, that activates endogenous RAS generating high levels of RAS-GTP, to quantify the upper bound on sensor output without RAS overexpression.

In order to quantify signal propagation along this cascade, we generated a number of reporter constructs. First, to quantify the expression of the RBDCRD-NarX fusion, we further fused this component to SYFP2 and measured YFP fluorescence in HEK293 cells with various RAS-GTP levels. We observed that overexpression of KRAS^G12D^ or overactivation of endogenous RAS with Sos-1 resulted in YFP increase and thus elevated expression of the RBDCRD-NarX fusion, compared to cells with baseline RAS-GTP. Conversely, inactivation of endogenous RAS with NF1 resulted in a lower YFP/RBDCRD-NarX expression (Fig. 2a). Second, we used the same construct to visualize RAS binding of the RBDCRD-NarX. Because RAS is a membrane protein^33^, we expected the binding of RBDCRD-NarX to KRAS-GTP to result in YFP accumulation at the membrane. Indeed, RAS-GTP increase by overexpressing KRAS^G12D^ or via Sos-1 led to higher membrane-to-total-signal ratio, compared to cells with endogenous RAS-GTP levels or with NF1-deactivated RAS (Fig. 2b). Third, we examined the dimerization of the RAS sensor by fusing the two parts of a split mVenus to RBDCRD-NarX proteins and measuring the signal from reconstituted mVenus. Again, RAS activation increased the signal from dimerized RBDCRD-NarX-mVenus, while RAS deactivation decreased it (Fig. 2c). Last, we investigated the effect of RAS-GTP levels on mCerulean sensor output of the complete RAS sensor. We observed the expected trend whereas the magnitude of the effect was amplified compared to the effects of individual sensing step, suggesting signal amplification in the synthetic sensing pathway (Fig. 2d).

**Figure 2.**
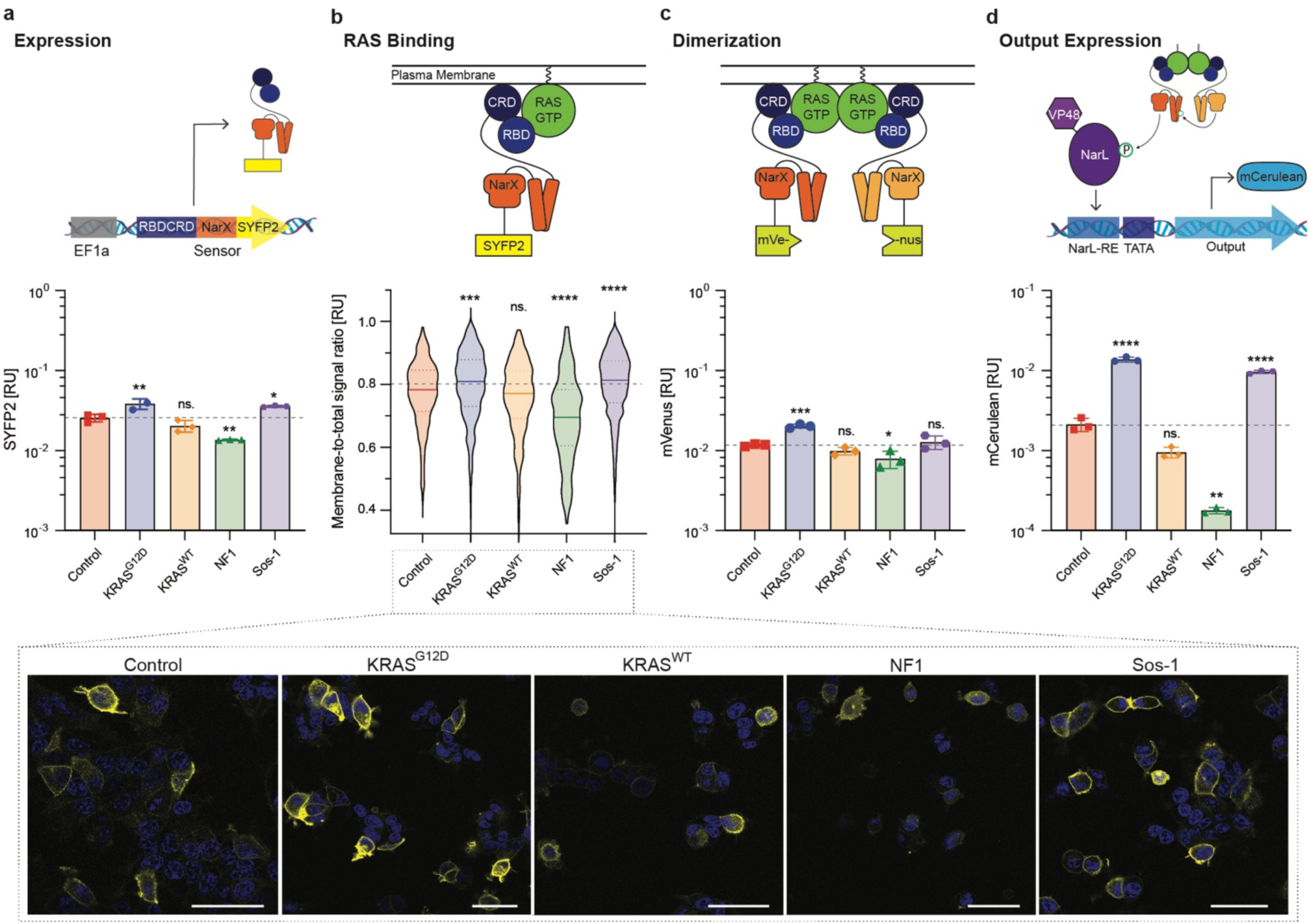
Mechanism of Action. Effect of differential RAS activation on the steps considered necessary for RAS Sensor activation. RAS activation in HEK293 cells was manipulated by co-expressing KRAS^G12D^, KRAS^WT^, Sos-1 (a guanine nucleotide exchange factor that activates endogenous RAS), or NF1 (a GTPase-activating protein that deactivates endogenous RAS). In the control condition the cells are transfected with a non-coding plasmid, here it represents the endogenous RAS activation. Schematics on top of the graphs illustrate how and what part of the Mechanism was investigated. (**a**) Expression levels of the RBDCRD-NarX-SYFP2 fusion protein measured by flow cytometry in the presence of various KRAS modulators (x axis labels). (**b**) RAS binding of the RBDCRD-NarX-SYFP2 fusion protein approximated by calculating the ratio of membrane to total SYFP2 signal for each cell. Intracellular localization of SYFP2 was measured using confocal microscopy. The micrographs below show representative images for each condition. Scale bars = 50 µm. (**c**) Dimerization of the NarX fusion proteins assessed by transfecting two complementary RBDCRD-NarX-split.mVenus fusions and measuring the mVenus fluorescence by flow cytometry. **(d**) Output expression after transfection with the full RAS Sensor measured by flow cytometry. In **a, c & d** the fluorescent signals were normalized to a constitutively expressed transfection control. Each symbol represents one biological replicate (**a**: n=9, **c-d**: n=3). The error bars represent +/- SD. In **b** the fluorescence at the membrane was normalized to the total fluorescence for each cell. Violin plots in **b** represent 560 (KRAS^G12D^), 322 (KRAS^WT^), 482 (Control), 226 (NF1), or 1194 (Sos-1) cells from three biological replicates. Significance was tested using an ordinary one-way ANOVA with Dunnett’s multiple comparisons to compare each condition with the control condition (endogenous RAS activation). *p < 0.05, **p < 0.01, ***p < 0.001, ****p < 0.0001.

While the dependency of RAS binding, dimerization, and output expression of the sensor on RAS-GTP levels was expected, we did not expect that the expression of the RBDCRD-NarX fusion itself (Fig. 2a) would be RAS-GTP dependent. Possible explanations could include the presence of transcription factor binding sites activated by RAS signaling in the elongation factor 1a (EF1a) promoter driving the expression of RBDCRD-NarX, or RAS-dependent non-specific activation of protein synthesis. EF1a promoter sequence contains potential binding sites of CREB, c-Myc, SRF, AP1 and Elk-4 (Supplementary Fig. 3)^34^. Additionally, there are reports that (over-)activated RAS or MAPK signaling can increase protein synthesis^35–38^ through post-transcriptional and translational mechanisms. These include upregulation of translation initiation factors and translational machinery^35^, as well as increased ribosome biogenesis^35,39^, suggesting that multiple mechanisms may be involved.

Comparing the functional RAS sensor to a panel of non-functional or constitutively active control sensors resulted in two additional insights: One, inactive controls lacking NarX dimerization show low increases in mCerulean output (Supplementary Fig. 4a), which are fully compensated –or slightly overcompensated– by normalization to the EF1a-driven mCherry transfection control (Supplementary Fig. 4b). Two, the RAS-dependent increase in expression is non-linearly amplified by the NarX-NarL system. Although the resulting 4-fold RAS-dependent increase in output signal is significantly lower than the 14-fold increase of the functional RAS sensor, suggesting that increased expression does not explain the full response (Supplementary Fig. 4b).

^2727^This indicates that dimerization and functional binding of RBDCRD to RAS are required for full sensor activation and output generation, which is further evidenced by the decreased activity of non-RAS binding mutants (Fig. 1e & Supplementary Fig. 4). While increased expression and its non-linear amplification are a contributing factor, RAS binding and RAS-dependent dimerization are necessary for achieving maximal dynamic range in RAS sensing.

### Optimization of activation efficiency and sensor tunability

The RAS sensor is activated by RAS-GTP and therefore by both wild-type and mutant RAS. Cells with mutant RAS have higher RAS-GTP and higher sensor activation than cells with wild-type RAS. Nevertheless, residual sensor activation in healthy cells with wild-type RAS may cause toxicity in a clinical implementation of the sensor. Therefore, we aimed to further improve the RAS sensor’s ability to discriminate between wild-type and mutant RAS and optimized the transfer function between RAS-GTP input and mCerulean output. Structural predictions with AlphaFold2^40^ motivated the exploration of longer linkers between the sensor’s RBDCRD and NarX-derived domains by showing that longer linkers may provide more flexibility for the NarX domains to get into close proximity which could improve dimerization and transphosphorylation efficiency (Fig. 3a). Our measurements showed that up to a certain size, longer and/or more rigid linkers indeed increased output expression, but not the dynamic range between KRAS^G12D^ and KRAS^WT^ (Fig. 3b).

**Figure 3.**
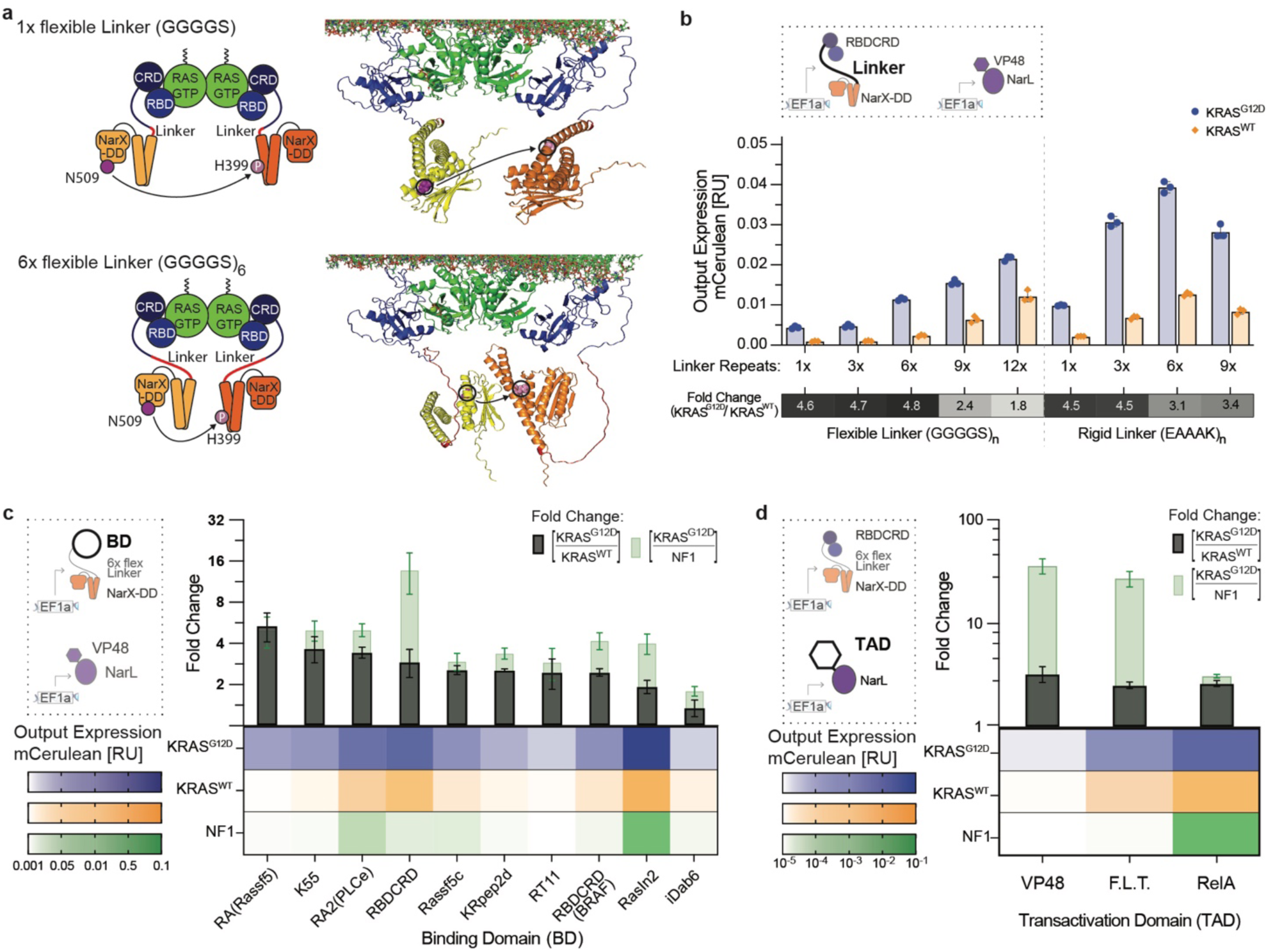
Tunability of the RAS sensor. (**a**) 3D structure of the RAS Sensor dimerizing at the membrane. The structure of the RBDCRD-NarX fusion proteins (orange and yellow) was predicted using AlphaFold and aligned with existing NMR structures of RBDCRD (blue) bound to a KRAS-dimer (green) at the membrane. The ATP-binding site N509 (purple) and the phosphorylation site H399 (pink) are highlighted as spheres. On the top RBDCRD and NarX are fused with a 1x GGGGS linker and on the bottom with a 6x GGGGS linker. (**b**) Effect of different linkers in the RBDCRD-NarX fusion protein of the RAS Sensor. The bar charts show the output expression in HEK293 co-transfected with KRAS^G12D^ (blue) or KRAS^WT^ (orange) when using different numbers of repeats of a flexible (GGGGS) or a rigid (EAAAK) linker in the RBDCRD-NarX fusion proteins. The heatmap below shows the corresponding fold change between output expression in cells with KRAS^G12D^ and KRAS^WT^. (**c**) Effect of different binding domains (BD) fused to NarX in the RAS Sensor. The heatmap shows the output expression in HEK293 co-transfected with 15 ng/well of KRAS^G12D^ (blue), KRAS^WT^ (orange), or NF1 (green), a GTPase-activating protein that deactivates endogenous RAS. The bars above show the corresponding fold changes between cells with KRAS^G12D^ and KRAS^WT^ (black) or KRAS^G12D^ and NF1 (green). (**d**) Effect of different transactivation domains (TAD) fused to NarL in the RAS Sensor. The heatmap shows the output expression in HEK293 co-transfected with KRAS^G12D^ (blue), KRAS^WT^ (yellow) or NF1 (green). The bars above show the corresponding fold changes between cells with KRAS^G12D^ and KRAS^WT^ (black) or KRAS^G12D^ and NF1 (green). mCerulean output expression was measured using flow cytometry and normalized to a constitutively expressed mCherry transfection control. Mean values were calculated from three (**b**) or two (**c-d**) biological replicates. Error bars were calculated using error propagation rules.

We also explored alternative natural and synthetic RAS binding domains. Among the tested binding domains, the Ras association domain (RA) of the natural RAS effector Rassf5, the RAS association domain 2 (RA2) of the phospholipase C epsilon (PLCe)^41^, and the synthetic RAS binder K55^42^ showed a slightly higher or similar dynamic range than RBDCRD (Fig. 3c). To better understand how different binding domains affect RAS-specific activation, we deactivated endogenous RAS^WT^ in HEK293 using NF-1. RBDCRD was the only binding domain that showed a pronounced decrease in output expression in cells with NF1 compared to cells with KRAS^WT^, indicating low RAS-unspecific activation for RBDCRD but also activation in cells with KRAS^WT^. The other binding domains showed one of two responses: (1) already very low activation in cells with KRAS^WT^ which was not further decreased in cells with NF1, for example in RA(Rassf5) and K55; (2) background activation in cells with NF-1 indicating RAS-unspecific activation, for example in RA2(PLCe) (Fig. 3c).

Fusing stronger transactivation domains to NarL markedly increased output expression of the RAS sensor without changing its dynamic range (Fig. 3d). Specifically, we replaced the initial VP48 with either F.L.T., a fusion of three partial transactivation domains, FoxoTAD (Forkhead-Box-Protein-O3^604-664^), LMSTEN (MYB^251-330^), and TA1 (RelA^521-331^) or with RelA^342-551^ with all its transactivation domains (Schweingruber et al. in preparation). In cells co-transfected with NF1, the sensor with NarL-F.L.T. exhibited low output expression similar to the initial sensor variant with NarL-VP48. In contrast, the sensor with NarL-RelA had high background in cells with NF1 suggesting RAS-unspecific activation when using RelA as a transactivation domain (Fig. 3d). Overall, the testing of alternative sensor parts did not increase the dynamic range between KRAS^G12D^ and KRAS^WT^ but it showed that the RAS sensor was modular. Different linkers, binding domains, and transactivation domains could be used to construct the sensor and tune the absolute strength of output expression without affecting the dynamic range.

### Multi-input RAS-targeting circuits with improved selectivity for KRAS^G12D^

Mutations in signaling proteins, such as RAS, often lead to increased activation of downstream transcription factors^43^. We hypothesized that using transcription factors from the mitogen-activated protein kinase pathway (MAPK) downstream of RAS as inputs could add an additional layer of selectivity to RAS-targeting circuits. Regulating the expression of RAS sensor components via MAPK-dependent response elements allowed to create a coherent type-1 feed-forward motif with AND-gate logic^44^ (Fig. 4a). This motif has been shown to act as a noise repressor^45^ and as a persistence detector with a delayed onset that is only activated by persisting input stimulation^44^. Both are desired properties in RAS-targeting circuits to detect constitutively active mutated RAS while minimizing output expression from the more transient activation of wild-type RAS^46,47^.

**Figure 4.**
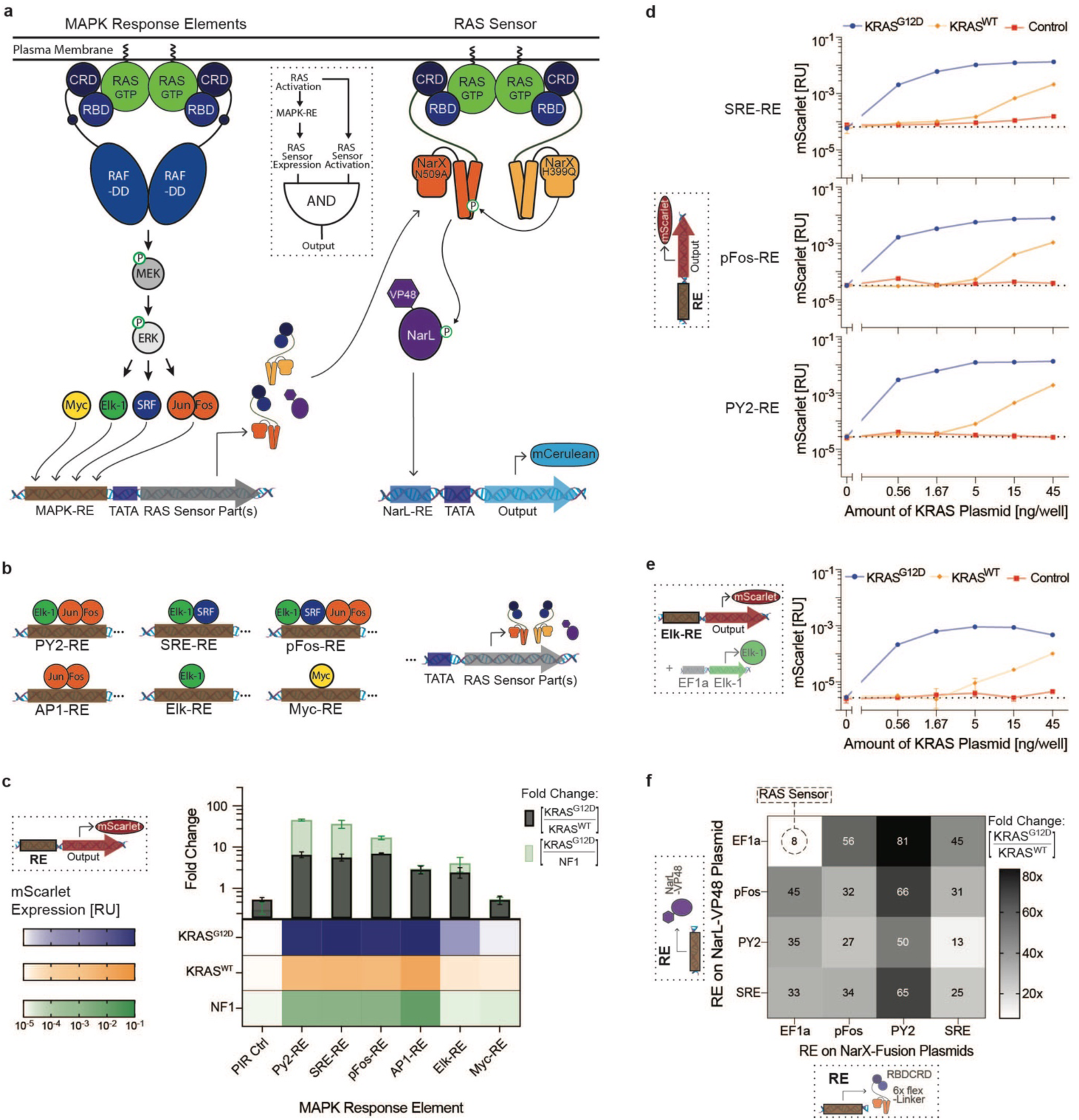
Design of Multi-Input RAS-targeting Circuits. (**a**) Schematic of the RAS-targeting circuit with an AND-gate between mitogen-activated protein kinase (MAPK) sensors and the direct RAS Sensor. Dimerization of RAS activates the MAPK pathway and its downstream transcription factors. These transcription factors bind the synthetic response elements (RE), expressing the parts of the RAS Sensor. The RBDCRD-NarX proteins then bind activated RAS, dimerize, and propagate the signal to NarL, leading to output expression. The logic diagram of the resulting coherent feed-forward loop with AND-gate logic is shown in the dotted box. (**b**) Schematic of the transcription factor binding sites present in the response elements. Multiple repeats of the binding sites were placed upstream of a minimal promoter (TATA) driving expression of the RAS Sensor parts. (**c**) Expression levels with the MAPK response elements. The heatmap shows the direct expression of mScarlet of the different REs in HEK293 cells co-transfected with 15 ng/well of either KRAS^G12D^ (blue), KRAS^WT^ (orange), or NF1, a protein that deactivates endogenous RAS (green). The bars above show the corresponding fold changes between cells with KRAS^G12D^ and KRAS^WT^ (black) or KRAS^G12D^ and NF1 (green). (**d**) RAS-dependency of the SRE-, pFos-, and PY2-response elements. Direct mScarlet expression of the response elements in HEK293 cells co-transfected with different amounts of KRAS^G12D^ (blue), KRAS^WT^ (orange), or non-coding control plasmids (red). (**e**) RAS-dependency of the Elk-RE when additionally overexpressing Elk-1. RAS titration as described in **d**. (**f**) Dynamic range of the RAS-targeting circuits. Fold change in mCerulean output expression between HEK293 co-transfected with 1.67 ng/well of KRAS^G12D^ and KRAS^WT^. In the RAS-targeting circuits, NarL-VP48 and/or the RBDCRD-6xfL-NarX fusion proteins were expressed using different MAPK-REs or a constitutive promoter (EF1a). Fluorescent protein expression was measured by flow cytometry and normalized to a constitutively expressed transfection control. Mean values were calculated from two (**c-e**) or three (**f**) biological replicates. PY2: polyoma virus enhancer domain; SRE: Serum response element; pFos: minimal promoter of c-fos; AP1: activator protein 1; Elk: Ets-like protein; Myc: myelocytomatosis protein. Detailed response element design is shown in Supplementary Fig. 5.

As an initial step, we designed synthetic response elements with binding sites for transcription factors shown to be activated by mutated RAS, including Elk-1^13^, c-Fos & c-Jun^48^, c-Myc^49^, and SRF^50^. For each response element, multiple binding sites of MAPK transcription factors^15,51,52^, minimal promoters from genes downstream of RAS^15^, or sequences from existing RAS-responsive promoters^16,22,53^ were encoded upstream of a low leakage minimal promoter^54^ (Fig. 4b & detailed in Supplementary Fig. 5).

To assess the functionality of the response elements, we cloned the response elements directly upstream of a fluorescent mScarlet protein and transfected them into HEK293 cells. The minimal promoter of c-fos (pFos)^15^, the polyomavirus enhancer domain (PY2)^22^, and the serum response element (SRE)^15^ were the response elements exhibiting the highest dynamic range between cells with KRAS^G12D^ and cells with KRAS^WT^ (Fig. 4c). The response elements demonstrated a similar dependency on KRAS^G12D^ and KRAS^WT^ to the one observed with the binding-based RAS sensor in Fig. 1e (Fig.4d). All three response elements contain binding sites for Ets-like 1 protein (elk-1). However, the Elk response element consisting of only elk binding sites showed low mScarlet expression and low dynamic range. To better understand this apparent contradiction, we further investigated the Elk response element. Overexpressing elk-1 with the Elk response reporter increased the mScarlet expression, indicating that endogenous elk-1 levels are not high enough for efficient activation of the Elk response element. With overexpressed elk-1, we observed high selectivity for cells with KRAS^G12D^ and low RAS-independent background activation in cells overexpressing NF-1 (Supplementary Fig. 6). A RAS titration provided more evidence for the RAS dependence of the Elk response element (Fig. 4e).

Superior performance of pFos, SRE, and PY2 response elements demonstrate that inclusion of additional transcription factor binding sites can amplify transcriptional sensor response, potentially due to synergy^55^. Consistent with this and previous reports that the combination of AP-1 and elk-1 binding sites is critical for maximal RAS responsiveness in the polyomavirus enhancer^16,56^, we found that combining elk-1 with additional transcription factors allows the design of efficient RAS-dependent response elements.

Finally, we combined these MAPK response elements with the RAS sensor using AND-gate logic, by using the response elements to regulate the expression of binding-triggered RBDCRD-NarX sensor. The resulting RAS-sensing circuits showed improved selectivity for KRAS^G12D^ with the best performer exhibiting an 81-fold dynamic range between KRAS^G12D^ and KRAS^WT^ compared to the 8-fold dynamic range of the binding-based RAS sensor alone (Fig. 4f). To assess the importance of RAS-binding in the RAS-targeting circuits, we replaced the RBDCRD-NarX fusion proteins with a constitutively dimerized non-truncated NarX control. The MAPK response element-expressed, non-truncated NarX showed dynamic ranges between KRAS^G12D^ and KRAS^WT^ from 2-fold to 20-fold. However, expressing RAS-binding dependent RBDCRD-NarX showed a higher dynamic range for all but the SRE_NarX + PY2_NarL combination (Supplementary Fig. 7). Thus, combining the MAPK response elements with the binding-based RAS sensor into RAS-targeting circuits generally improved the distinction between cells with KRAS^G12D^ and KRAS^WT^ and allowed to reach higher maximal fold changes.

### Modularity allows to create RAS sensor circuits with high dynamic range

As shown above, RAS-sensing circuits can utilize a variety of MAPK response elements, RAS binding domains, linkers, and transactivator domains. To further optimize the transfer function between the input and the output and improve the discrimination between cells with mutant and wild-type RAS, we performed a screening campaign to examine the effect of different combinations of circuit components on the dynamic range. In this screen we tested combinations of various building blocks: RBDCRD, K55, and RA(Rassf5) as binding domains fused to the NarX; VP48 or F.L.T as transactivation domain of NarL; and pFos, PY2 and SRE as MAPK response elements. The response elements regulated the expression of either the NarX fusion proteins only, or of the NarL protein only, while the other part was constitutively driven by EF1a. For the EF1a-driven RBDCRD-NarX proteins we tested two versions with either a 6x flexible or a 3x rigid linker. This set resulted in a total of 222 unique conditions, over two experimental batches. Almost thirty conditions resulted in high dynamic ranges of >100-fold, with the highest circuit variants using the combination of PY2, RBDCRD, and F.L.T. (Fig. 5a).

**Figure 5.**
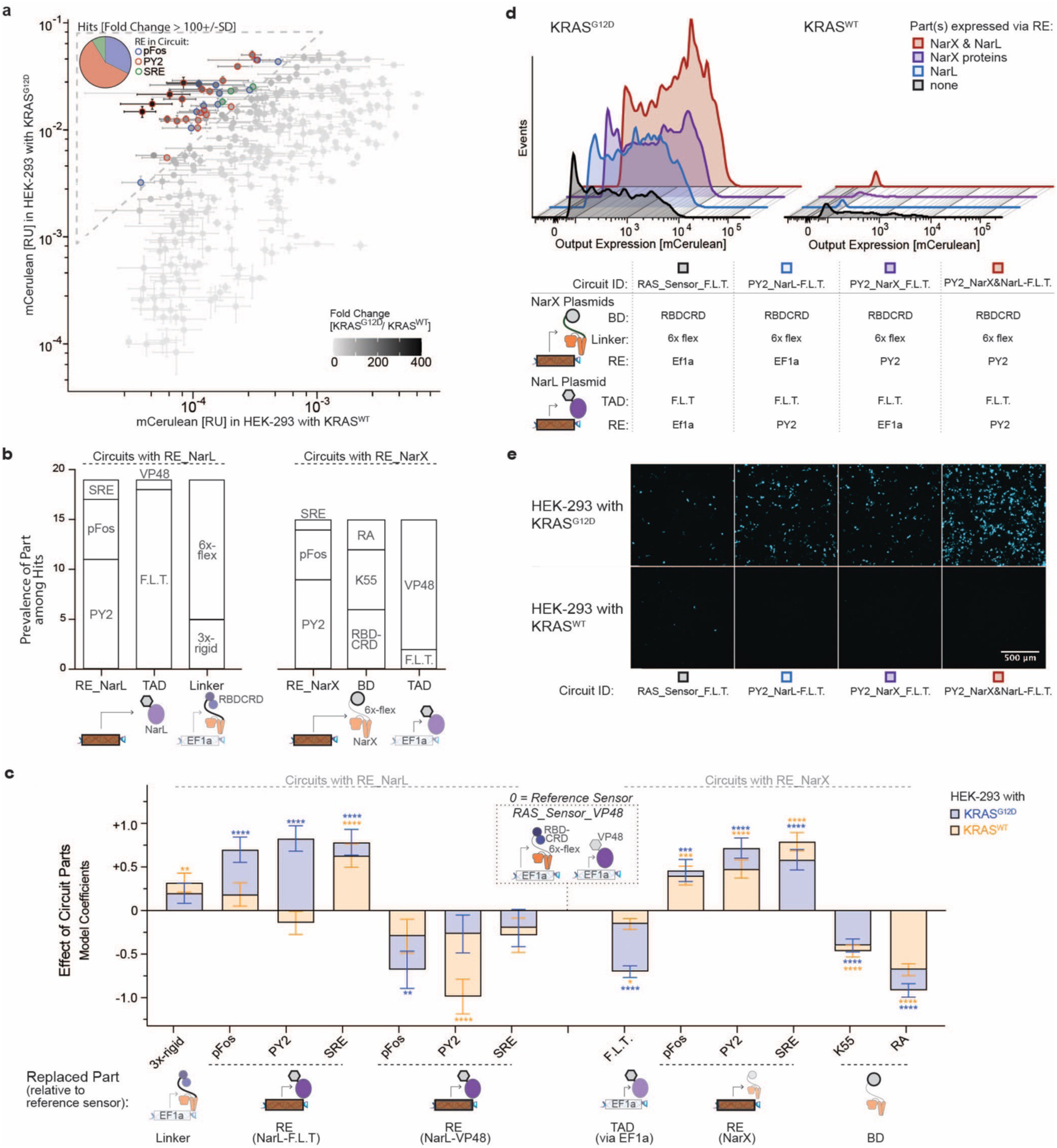
Modularity of the RAS-targeting Circuits. (**a**) Screening of RAS circuit variants with different parts. mCerulean output expression in HEK293 co-transfected with 1.67 ng/well of KRAS^G12D^ versus KRAS^WT^ representing the ON-versus OFF-state of the screened circuits. The gray shading of the symbols represents the dynamic range (fold change between ON and OFF). Circuits with a high dynamic range (> 100+/-SD) are highlighted. The pie chart shows the prevalence of the response elements (RE) among the hits. (**b**) Prevalence of the circuit parts among the hits. On the left for all circuits where the RE expressed NarL (RE_NarL) and on the right where the RE expressed the NarX proteins (RE_NarX). TAD: transactivation domain, BD: binding domain. RA: Ras association domain of Rassf5 (**c**) Effect of different circuit parts on the output expression in HEK293 with KRAS^G12D^ (blue) and KRAS^WT^ (orange) fitted using a generalized linear regression. The EF1a expressed RAS Sensor with RBDCRD, a 6x flexible linker in the NarX fusion proteins, and VP48 as TAD fused to NarL was set as the reference sensor. The graph shows the model coefficients, which can be interpreted as the effect on output expression when a part in the reference sensor is replaced by the part indicated on the x-axis. (**d**) Expression of all circuit parts via response elements. Fluorescence histograms of mCerulean-positive cells comparing ON-(KRAS^G12D^) and OFF-state (KRAS^WT^) of RAS circuits when either NarL (blue), the NarX proteins (violet), or all parts (red) are expressed via REs. The tested circuits contain RBDCRD as BD, a 6x flexible linker and F.L.T. as TAD, the parts that led to the hits with the highest dynamic range. The table below shows the parts used in each of the tested circuit. (**e**) Microscopy images showing the mCerulean expression of the conditions from **d**. mCerulean output expression was measured by flow cytometry and normalized to a constitutively expressed mCherry transfection control. Each circuit was measured in three biological replicates. Error bars represent +/- SD. Significance in **c** was tested using the Wald test. *p < 0.05, **p < 0.01, ***p < 0.001, ****p < 0.0001.

We found that PY2 was the most prevalent response element among the high-performing hits, followed by pFos, while SRE only occurred in three hits. The prevalence of the transactivation domain varied depending on whether the NarX or the NarL proteins were regulated by a MAPK response element. While F.L.T. was more abundant among the hits where NarL was regulated by the response elements, VP48 dominated in the circuits where the response elements regulated the NarX fusion proteins. The K55 and RBDCRD binding domains were found in an equal number of hits (Fig. 5b). However, we observed that circuits with F.L.T. as the transactivation domain performed best with RBDCRD as the binding domain, while there was no significant difference between RBDCRD and K55 for circuits with VP48 (Supplementary Fig. 8).

To further improve our understanding of the effect of different building blocks, we fitted a linear regression model (Fig. 5c) and compared it with the experimental data (Supplementary Fig. 9-10). This revealed that the effect of the MAPK response elements depended on which sensor components they regulated. Expression of NarL-F.L.T. under the control of MAPK response elements strongly increased the output expression in cells with KRAS^G12D^, while, except for the SRE_NarL-F.L.T. circuit, the activation in cells with KRAS^WT^ remained low. In contrast, MAPK control of NarL-VP48 decreased the output in KRAS^G12D^ and even more so in KRAS^WT^ cells. Expression of the NarX fusion proteins increased the circuit activation in cells with KRAS^G12D^, but also in cells with KRAS^WT^. (Fig. 5c, & Supplementary Fig. 9a-f).

In this campaign, we focused on circuits where only NarL or only the NarX proteins were regulated by the response elements. Next, we also implemented the regulation of all circuit components by PY2. The PY2_NarX & NarL-F.L.T. circuit led to the highest output in cells with KRAS^G12D^, with only marginally higher activation with KRAS^WT^ than PY2_NarX_F.L.T. or PY2_NarL-F.L.T. and still lower than the RAS_Sensor_F.L.T. (Fig. 5d&e).

In summary, the tests revealed that the RAS-targeting circuits are modular, with various combinations of parts leading to circuits with high dynamic ranges that diverge in their transfer functions and their activation thresholds. The availability of different circuit parts thus allowed to adapt the input-output behavior of the circuits and increased the distinction between cells with mutated and cells with wild-type RAS.

### RAS sensor circuits detect endogenous RAS mutations in cancer cells

To evaluate the response of the RAS sensor circuits to endogenous RAS activation in cancer cell line, we transfected the EF1a-driven, binding-triggered RAS sensor into wild-type HCT-116 cells, a colon cancer cell line harboring a KRAS^G13D^ mutation (HCT-116^WT^)^57^. To have a comparable off-target cell line without mutated RAS, we also transfected a commercially available KRAS knock-out HCT-116 cell line (HCT-116^k.o.^).

The RAS sensor was functional in HCT-116 cells and responded to endogenous levels of RAS activation with higher activation in HCT-116 cells with KRAS^G13D^ than in the knock-out cells. Further, loss-of-function mutations in RBDCRD decreased activation (Fig. 6a). However, the dynamic range was only 3-fold (Supplementary Fig. 11). Therefore, we leveraged the modularity of the circuit design to improve selectivity for target HCT-116 cells. We found that the amplified AND-gate circuits with PY2 response element, F.L.T transactivation domain, and RBDCRD as a RAS-binding domain, were most effective in distinguishing HCT-116^WT^ from HCT-116^k.o.^ (Fig. 6b-d). In contrast to what we saw in HEK293 overexpressing RAS (Fig. 5d), the “AND-gate” RAS-targeting circuits do not generate higher output than the EF1a-driven, binding-triggered RAS sensor in HCT-116. Instead, the improved dynamic range results from decreased leakiness in HCT-116^k.o.^. Only with a 3x rigid linker in RBDCRD-NarX fusion did the PY2_NarL-F.L.T circuit show higher output expression compared to the EF1a-driven, binding-triggered RAS sensor. However, while the circuit with the 3x rigid linker still showed a dynamic range of 18-fold it had a decreased dynamic range compared to the same circuit with the 6x flexible linker (57-fold) (Fig. 6e). Taken together, this dataset demonstrates that the RAS-targeting circuits are functional in cancer cells and are able to respond to endogenous levels of mutated RAS. While there are differences between the model systems, we can leverage the availability of multiple circuit parts to adapt the circuits to specific target and off-target cells to improve selectivity.

**Figure 6.**
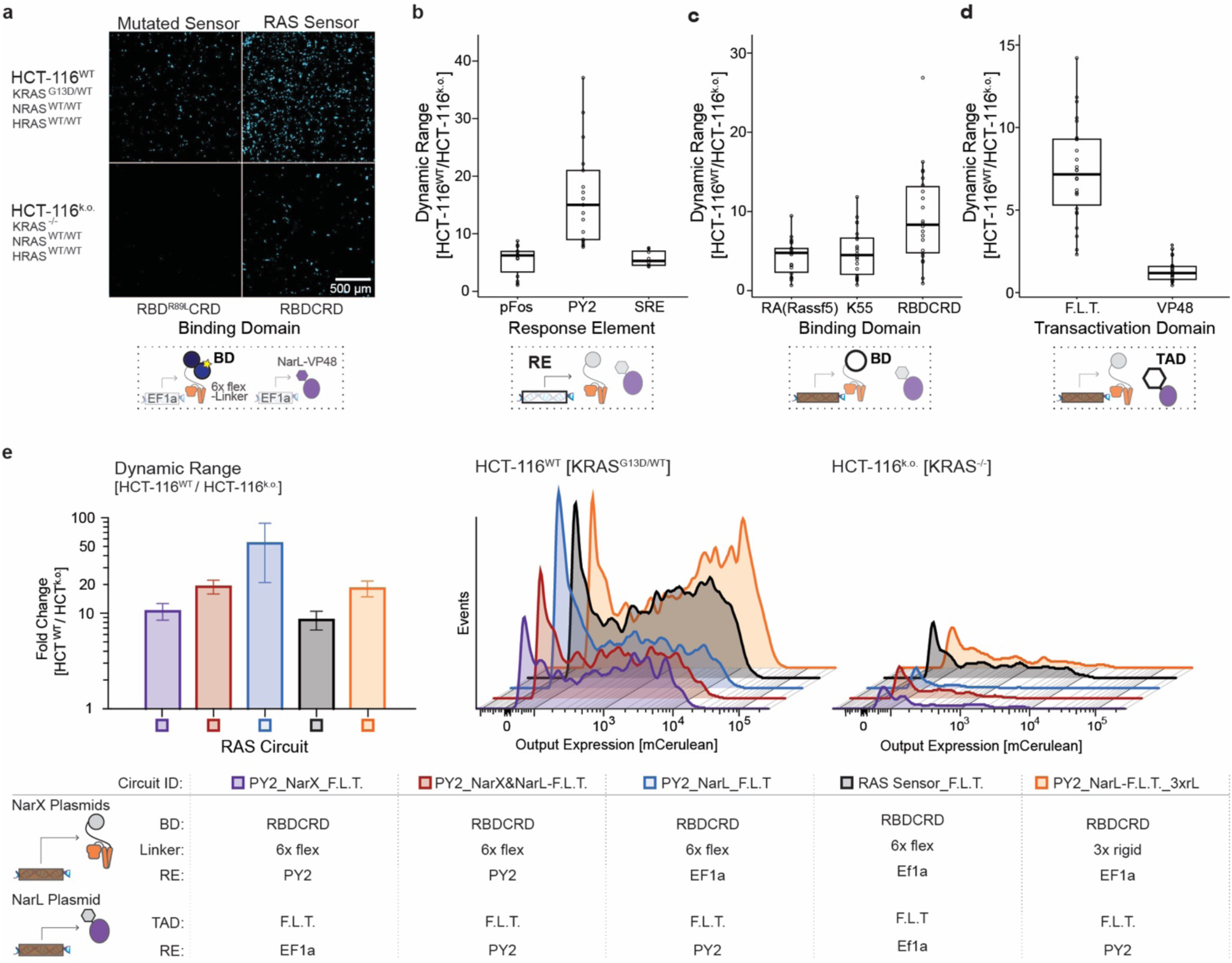
Translation into cancer cells – Detection of endogenous RAS levels in HCT-116. (**a**) RAS Sensor activation in colon cancer cells. Microscopy images of the mCerulean output expression in HCT-116 wild-type cells harboring a homozygous KRAS^G13D^ mutation (HCT-116^WT^; top row) and HCT-116 KRAS knock-out cells (HCT-116^k.o.^; bottom row) transfected with the initial RAS Sensor (right) or a RAS Sensor with an R89L mutation in the Ras binding domain (left). (**b-d**) Effect of different circuit parts in colon cancer cells. Boxplots of dynamic range of different RAS-targeting circuits grouped by the circuit parts of interest they contain. The circuit parts investigated were: the response elements in **b**, the binding domain fused to the NarX proteins in **c**, and the transactivation domain fused to NarL in **d**. Each black circle represents a different RAS circuit. (**e**) Best performing RAS-targeting circuits in colon cancer cells. The parts used in each RAS circuit are listed in the table below. The bar graph shows the dynamic range, while the fluorescence histograms show mCerulean-positive cells obtained in the On-(HCT-116^WT^) and Off-state (HCT-116^k.o.^) of the circuits. mCerulean output expression was measured by flow cytometry and normalized to a constitutively expressed mCherry transfection control. Dynamic range was calculated as fold change between normalized output expression in HCT-116^WT^ and HCT-116^k.o.^. Each circuit was measured in three biological replicates. Error bars in **e** were calculated using error propagation rules.

### RAS circuits can generate selective output in RAS-driven cancer cells

To specifically target RAS-driven cancer, the RAS-targeting circuits need to show selective output expression in cancer cells with mutations over-activating RAS (RAS^MUT^), while maintaining minimal expression in cells without such mutations (RAS^WT^). Testing the most promising RAS-targeting circuits in twelve cancer cell lines showed that all circuits exhibited significantly higher output expression in RAS^MUT^ cells (Fig. 7a). The PY2_NarX&NarL-F.L.T. circuit had the highest response rate among the RAS^MUT^ cells but displayed slightly increased background activation in RAS^WT^ cells. This was particularly notable in HT-29, a cell line harboring a BRAF mutation^57^ and thus representing a RAS^WT^ cell line but with an over-activated MAPK pathway. HT-29 did not show an elevated background in any of the other circuits, indicating a clean AND-gate behavior between the RAS Sensor and the MAPK response element for the circuits that express only NarL-F.L.T. with the response elements (Fig. 7a).

**Figure 7.**
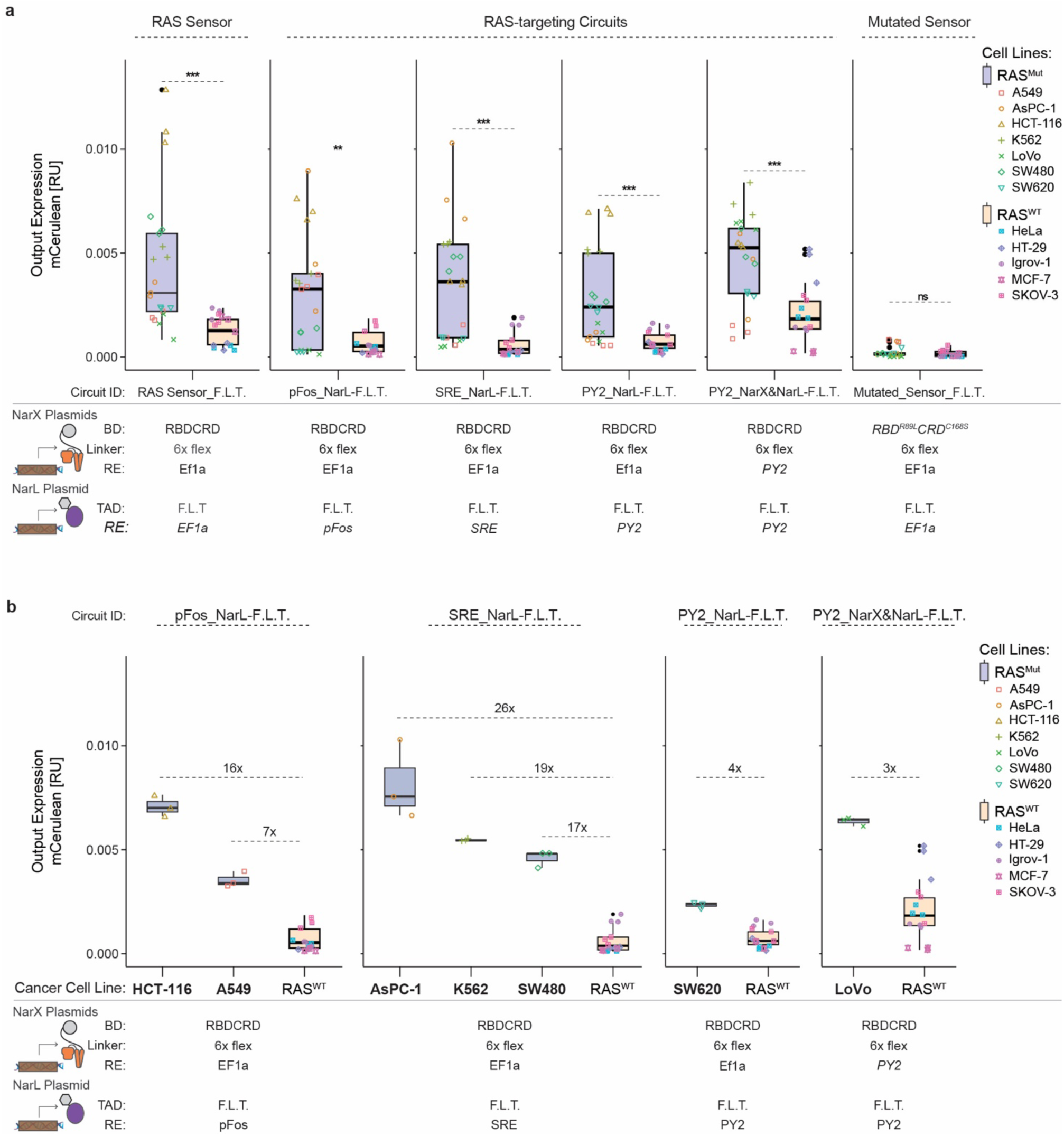
Translation into cancer cells – Selectivity for RAS-driven cancer cells. (**a**) RAS-targeting circuits are classifiers for cells with mutated RAS. Output expression of RAS-targeting circuits in different cancer cell lines with (RAS^MUT^ = blue) or without (RAS^WT^ = orange) mutation leading to increased RAS activation. (**b**) Output expression of the best performing RAS-targeting circuits for each RAS^MUT^ cancer cell line. Each RAS^MUT^ cell lines is only shown in the circuit that performed best in the respective cell line. A boxplot of all RAS^WT^ cell lines is shown in each circuit and fold changes were calculated between the mean of the individual RAS^MUT^ and the mean of all RAS^WT^ cell lines. The colored symbols represent biological replicates of the different cell lines. The parts used in each RAS circuit are indicated in the tables below. mCerulean output expression was measured by flow cytometry and normalized to a constitutively expressed mCherry transfection control. Each circuit was measured in three biological replicates. Significance was tested using an unpaired two-tailed Student’s t-test. **p < 0.01, ***p < 0.001.

The output expression of the RAS circuits correlated with the direct expression of a fluorescent protein via the MAPK response elements, although not perfectly (Supplementary Fig. 12). This indicates that, while not the only factor, the response elements are important for the differential activation between the cell lines.

Changing the response element that regulates NarL-F.L.T. expression in the RAS circuit, altered which RAS^MUT^ cells showed output expression. While some RAS^MUT^ cell lines, such as HCT-116 and K562, had higher output expression than all RAS^WT^ cells with all tested circuits, other RAS^MUT^ cell lines only showed higher expression with certain circuits, indicating that not all response elements work equally well in all RAS-driven cancer cell lines (Fig. 7a & Supplementary Fig. 13). The availability of different response elements enabled us to identify for each RAS^MUT^ cell line a functional RAS circuit with higher activation in these cells than in all RAS^WT^ cell lines (Fig. 7b & Supplementary Fig. 13). For example, while the PY2_NarL-F.L.T. circuit did not show higher activation in AsPC-1 than in the RAS^WT^ cell lines, using SRE instead of PY2 led to a 26-fold higher output expression than in the RAS^WT^ cell lines (Supplementary Fig. 8 & Fig. 7b). This demonstrates that the response elements allow to adapt the RAS circuits to the targeted cancer cell type.

### RAS circuits can kill RAS-driven cancer cells

To bring RAS-targeting circuits closer to therapeutic application, we replaced the fluorescent reporter with a clinically relevant output protein: a herpes simplex virus thymidine kinase variant (HSV-TK)^58^. HSV-TK functions as a suicide gene by converting the non-toxic prodrug ganciclovir (GCV) into a cytotoxic triphosphate derivative^21^.

RAS-targeting circuits expressing HSV-TK induced robust cell death in KRAS-mutated HCT-116 cells after treatment with 50 µM GCV (Fig. 8a). Compared to the non-toxic GFP-output control (GFP-circuit) condition, where cells reached full confluence, EF1a_RAS-Sensor_F.L.T. and PY2_NarL-F.L.T. reduced confluence at 180h 1.8- and 1.7-fold, respectively. The positive control (EF1a_HSV-TK) reduced confluence 2.9-fold. Looking only at transfected cells, the effect was even more pronounced with EF1a_RAS-Sensor_F.L.T. and PY2_NarL-F.L.T. showing a 2- and 2.6-fold lower fluorescence than the GFP-circuit, respectively (Fig. 8b). Microscopy at 180h confirmed substantial killing. While GFP-circuit wells showed dense monolayers, those transfected with RAS-targeting circuits or EF1a_HSV-TK contained debris and rounded, aggregated cells, indicating low viability (Fig. 8c). These results show that HSV-TK enables the RAS-targeting circuits to efficiently kill RAS-mutant HCT-116 cells, including non-transfected neighbors likely via a bystander effect^59^.

**Figure 8.**
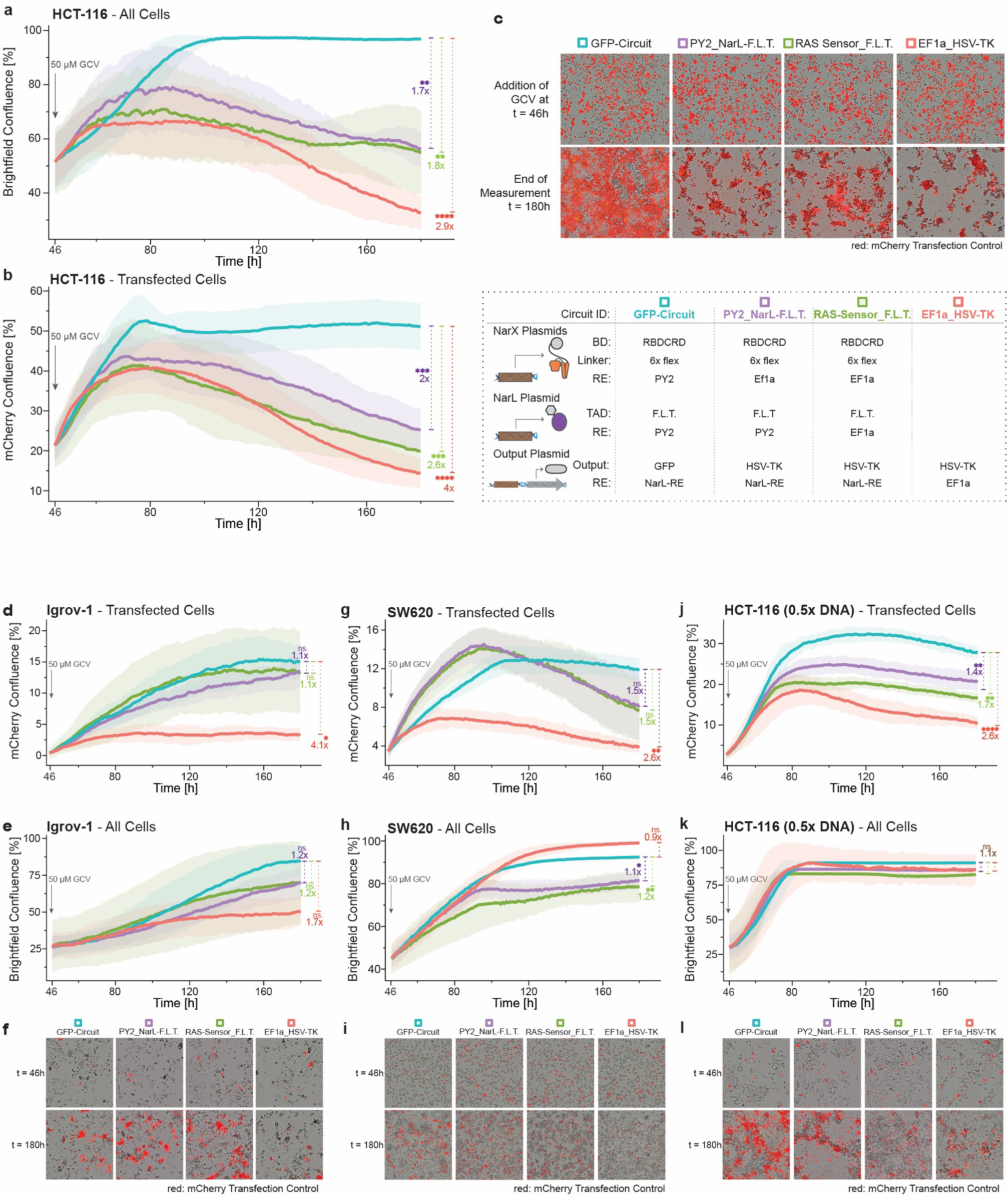
Killing of RAS-driven cancer cells. The graphs show the overall confluence, or confluence of mCherry transfection control positive cancer cells, transfected with RAS-targeting circuits that express herpes simplex virus thymidine kinase (HSV-TK) as output protein or controls over time. The used RAS circuits (purple and green) and controls are described in the dotted box, with a RAS circuit expressing GFP as output as negative control without HSV-TK (turquoise) and a EF1a-expressed constitutive HSV-TK as positive control (red). Gray arrow at t=46h indicate the addition of the prodrug ganciclovir (GCV) that is activated by HSV-TK. Statistical significance and fold chances are calculated between each condition and the corresponding GFP-circuit control at t=180h. Representative microscopy images of both brightfield and mCherry confluence at the time of ganciclovir addition (t=46) and end of the measurement (t=180) are shown for each condition and cell line. (**a**) Overall confluence in the well of KRAS^G13D^-mutated HCT-116 cells. (**b**) Confluence of transfected KRAS^G13D^-mutated HCT-116 cells. (**c)** Microscopy images corresponding to a & b. (**d**) Overall confluence in the well of RAS wild-type Igrov-1 cells. (**e**) Confluence of transfected RAS wild-type Igrov-1. (**f)** Microscopy images corresponding to d & e. (**g**) Overall confluence in the well of KRAS^G12V^-mutated SW620 cells. (**h**) Confluence of transfected KRAS^G12V^-mutated SW620 cells. (**i)** Microscopy images corresponding to g & h. (**j**) Overall confluence in the well of KRAS^G13D^-mutated HCT-116 cells transfected with lower (0.5x) amounts of the circuits. (**k**) Confluence of transfection control positive KRAS^G13D^-mutated HCT-116 cells transfected with lower (0.5x) amounts of the circuits. (**l)** Microscopy images corresponding to j & k. Confluence was quantified from microscopy images. Mean confluence was calculated from biological triplicates and background adjusted to the EF1a_HSV-TK condition of the same cell line at t=46h. Standard deviation is shown as ribbons. Significance was tested using an ordinary one-way ANOVA with Dunnett’s multiple comparison test. ns = non-significant, *p < 0.05, **p < 0.01, ***p < 0.001, ****p < 0.0001.

Next, we tested the circuits in Igrov-1, the RAS wild-type line that previously showed the highest activation among RAS^WT^ cells for both circuits (Fig. 7a). RAS-targeting circuits with HSV-TK did not prevent growth in Igrov-1. Following 50 µM GCV treatment, both transfected and total cell populations continued to grow (Fig. 8d&e). In contrast, constitutive HSV-TK expression inhibited growth of transfected cells (Fig. 8d), although without a significant effect on bystander cells (Fig. 8e), likely due to lower transfection efficiency than HCT-116. Compared to the non-toxic GFP-circuit, EF1a_HSV-TK caused a 4-fold reduction in confluence of transfected cells, while both RAS circuits showed a non-significant, 1.1-fold reduction of growth (Fig. 8d & microscopy in Fig. 8f). This suggests low toxicity of the RAS circuits in these RAS^WT^ cells. However, it is important to note the lower transfection efficiency of Igrov-1 (Fig 8d), compared to HCT-116 (Fig. 8a).

To assess the effect of transfection efficiency, we tested SW620 –a RAS^MUT^ cell line with lower transfection efficiency than HCT-116– and HCT-116 transfected with only half the DNA dose. Lower circuit dose or transfection efficiency also showed killing of RAS-driven cancer cells, but a less pronounced effect on bystander cells (Fig. 8g-l). SW620 and Igrov-1 had similar transfection efficiency (Supplementary Fig. 14a) but distinct killing curves with SW620 showing a decrease and Igrov-1 showing an increase in confluence of transfect cells (Fig. 8d & 8g), supporting the notion of selective cytotoxicity in cells with mutated RAS. However, the differences observed in transfection efficiency (Supplementary Fig. 14a–b), growth characteristics (Fig. 14c–d), and GCV/Ef1a_HSV-TK-sensitivity (Fig. 14e–f) may also influence RAS circuit-mediated killing, indicating that comparisons across cell lines should be interpreted cautiously.

In summary, RAS-targeting circuits expressing HSV-TK induced cell death in transfected RAS-mutant lines (HCT-116 and SW620), while RAS wild-type Igrov-1 cells continued to grow. Although cell-line-specific differences limit final conclusions about selectivity, our data supports a preferential cytotoxic effect in RAS-mutant cells. Remarkably, in KRAS^G13D^-mutated HCT-116 cells, higher DNA doses led to near-complete eradication of both transfected and neighboring cells, validating the potential of RAS-targeting circuits to express clinically relevant output proteins and effectively kill RAS-driven cancer cells.

## Discussion

In this study, we report the design of synthetic gene circuits to target RAS-driven cancers. We developed a set of RAS sensors with interchangeable parts that can be combined to flexibly design RAS-sensing circuits. Using our modular design, we created and characterized gene circuits advancing RAS-targeting circuits on two key performance criteria: selectivity for mutated RAS, and adaptability to different target and off-target cell lines.

To date the only existing synthetic gene circuit that directly targets RAS had a 2-fold dynamic range between HEK cells overexpressing KRAS^G12V^ and KRAS^WT^ and a 4-fold dynamic range when compared to HEK cells with endogenous RAS levels^24^. Our design of RAS-targeting circuit greatly improves specificity for mutant RAS. This is critical given that RAS circuits sense activated RAS-GTP, which is highly overactivated in cancers with RAS mutations^25^ but also present in healthy cells with wild-type RAS. Since this could lead to on-target, off-tumor effects, RAS-targeting circuits are designed to sense the different activation dynamics and activation levels^46,47^ resulting from the constitutive overactivation of RAS in cancer^25,60^. To achieve high dynamic range between cells with mutated and cells with wild-type RAS, we optimized the transfer function governing the relationship between the RAS-GTP input and sensor output. To this end, we combined binding-triggered RAS sensors and RAS-dependent MAPK transcription factor sensors into a coherent type 1 feed-forward AND-gate. In this design, MAPK sensors enhance the RAS-dependent increase in expression of the binding-triggered RAS sensor components. Thus, cells with wild-type RAS have lower levels of the RAS sensor components than cells with mutant RAS, leading to suppression of RAS-dependent leakage of output production by the binding-triggered RAS sensor. This network motif was shown to delay the onset of output expression but not the output shutdown^44^. This dynamic behavior, termed sign-sensitive delay ^44^, acts as a noise repressor ^45^ and persistence detector ^44^, which could further explain why this design improves the sensing of constitutively active mutant RAS while minimizing output expression from the more transient activation of wild-type RAS^46,47^. Integrating multiple RAS dependent sensors into the circuit design resulted in a strongly enhanced distinction between cells overexpressing mutant or wildtype RAS and a more than 100-fold higher dynamic range than previous circuits^24^.

Validation in cancer cell lines demonstrated that the RAS-targeting can selectively express an output in cancer cells. The RAS circuits are activated by all three RAS isoforms and a variety of mutations which allowed targeting of cancer cells with diverse KRAS mutations, but also of K562 cells with a BCR-ABL fusion gene that constitutively activates RAS through Sos-1^61^. A broad targeting range is advantageous because tumors exhibit high intra- and intertumoral heterogeneity of RAS mutations^62^. While broader RAS inhibitors are under development^8^, RAS heterogeneity is a limiting factor in current KRAS^G12C^ inhibitors because of resistance development due to escape variants with different RAS mutations^5^. The broad target range of the RAS-targeting circuits enabled us to identify a circuit for each mutant RAS cell line that was more active in those cells than in all wild-type RAS cell lines. However, the heterogeneous cancer cell lines showed variable dynamic ranges, indicating a need for adaptation of the circuits to the target cells.

Additional input sensors may allow to further enhance the RAS-targeting circuits. Beyond the direct RAS sensors and the MAPK response elements, adding other RAS-dependent sensors, such as ERK sensors^63,64^ or RAS-dysregulated miRNAs^13,65^ in an AND-gate configuration could further improve specificity. Alternatively, an OR-gate configuration could reduce the risk of resistance development. Off-target effects could be minimized by including NOT-gates with inputs associated with healthy cells, such as existing p53 sensors^17,66^ or new sensors specific for healthy cells with high RAS activation^67,68^.

In the context of cancer therapies, synthetic gene circuits can express a variety of therapeutic proteins as output, such as proapoptotic proteins^18,23^, enzymes that convert a prodrug into a cytotoxic drug^21^, immunotherapeutic proteins^53^, or even combination therapies^53^. We demonstrated that exchanging the output protein can be achieved by changing the coding sequence on the output plasmid. Armed with a therapeutic output, such as HSV-TK, RAS-targeting circuits can robustly kill the RAS-driven cancer cell lines HCT-116 and SW620. Simultaneously, these RAS circuits did not prevent growth in Igrov-1, suggesting low toxicity in this wild-type RAS cell line. While this indicates preferential cytotoxicity in RAS-driven cancer cells, cell line specific differences, such as in transfection efficiency, limit conclusions, underscoring the importance of further validation.

This further highlights another remaining challenge: delivery. The multi-plasmid-based delivery is likely difficult to implement in RAS-driven solid cancers. Thus, future efforts should aim at integrating all components on a single vector. For DNA, viral delivery is generally most efficient but has limited packaging capacity^69^. Compared to EF1a, the MAPK response elements reduce the overall size of the constructs to approximately 5 kb, bringing them well within the 8 kb packaging capacity of lentiviral vectors^69^ and leaving space for outputs, such as the 1.5 kb NarL-RE_HSV-TK cassette. The assembly of multiple modules on a single vector provides challenges for synthetic gene circuits, including positive and negative interactions between promoters and genes in proximity^70^. Using the MAPK-driven circuit versions without constitutive promoters may provide some robustness. Unintended direct output expression by neighboring MAPK response elements would still retain a certain RAS dependency, reducing the risk of constitutive, non-specific output expression in healthy cells. Nonetheless, assembling the circuit on a single vector will require careful design and rigorous validation to ensure optimal performance.

Reaching every single cancer cell will be challenging with any delivery system. We have seen that HSV-TK can also kill non-transfected, neighboring cancer cells suggesting that outputs with bystander effect are potentially more effective. However, this effect was strongly dose-dependent, and it may require precise dosing to optimize the killing of cancer cells while minimizing toxicity. In addition to dosing the DNA amount, selection of the components used in our RAS circuits tunes the expression strength, which may allow further dosing of HSV-TK but also adaptation when using different therapeutic output proteins. Potent molecules, such as interleukin-12, will require stringent expression with low leakiness to prevent systemic toxicity^4^, while less toxic molecules could benefit from stronger expression. We envision that the modularity of our system will allow to tailor the expression profile to the therapeutic output. In conclusion, this study provides the foundation for the design of RAS-targeting circuits. Our results confirm the feasibility of developing synthetic gene circuits that selectively target RAS-mutated cancer cells, demonstrate robust killing of certain RAS-driven cancer cells, and encourage the use of their versatility to adapt the circuits to future challenges during clinical translation. *While alternative delivery systems and validation in more realistic models will be essential*, this highlights the potential of RAS-targeting circuits as a new therapeutic strategy that could reshape therapies against RAS-driven cancer

## Methods

### Plasmid Construction

Plasmids were cloned using standard cloning techniques, such as Gibson, GoldenGate assembly, or restriction enzyme cloning. DNA fragments were ordered from TWIST Biosciences or Genewiz (Azenta Life Sciences). Enzymes were purchased from New England Biolabs and Thermo Fischer Scientific. The sequences of all used plasmids are listed in Supplementary Table 1 and the sequences of the individual RAS circuit parts in Supplementary Table 2.

### Cell Culture

All cell lines used were cultured at 37 °C, 5% CO2 in the medium suggested by the provider with 10% FBS and 1% penicillin/streptomycin solution. Cells were passaged before reaching confluency 1-3 times per week, depending on experimental plans. Details on the individual cell lines, such as provider, medium, and splitting ratios are listed in Supplementary Table 3.

### Transfections

The transfections for the RAS-GTP pulldown experiment in Fig. 1f and the confocal microscopy experiment in Fig. 2b were performed in 6-well plates, the transfection for the microscopy experiment in Fig. 6a, and the cell line screening in Fig. 7 in 24-well plates, and all other experiments were performed in 96-well plates. The difference in plate size was accounted for by scaling the amount of transfected DNA to the number of seeded cells. The cells were seeded 24h before transfection in 100 µL of medium (500 µL for 24-well; 2.5 mL for 6 well plates). Endotoxin-free (ZR DNA Prep Kit, Zymo Research, cat.no. D4201 & D4212) plasmids were mixed according to the experimental layout (Supplementary Tables 4 & 5), Opti-MEM^TM^ (Thermo Fischer Scientific) was added to reach a volume of 30 µL (50 µL in 24-well plates; 500 µL in 6-well plates). Transfection reagents were mixed with Opti-MEM^TM^ to reach a volume of 20 µL (50 µL in 24-well plates; 500 µL in 6-well plates). After incubation for at least 5 minutes, the mixture was added to the DNA samples, gently vortexed, spun down, and then incubated at room temperature for at least 20 minutes before addition to the cells. The seeding and transfection conditions for each cell line are shown in Supplementary Table 6 for the cell line screening in Fig. 7 and in Supplementary Table 7 for all other experiments. To minimize the effect of differential evaporation in 96-well plates, only the inner 60 wells were used for samples, while the outer wells were filled with PBS.

For the experiments comparing different cell lines (Fig. 6, 7 & 8d-l & Supplementary Fig. 8, 9, 13 & 14), the DNA amount was optimized to achieve more similar transfection efficiencies, as listed in Supplementary Tables 6-7. In the cancer cell line screening, we adjusted the DNA amount to 1.5x for cell lines with low transfection efficiencies (<20%), 1x DNA for cell lines with moderate efficiency (20-50%), and 0.5x DNA for cell lines with high efficiency (>50%).

### Structure Prediction

The protein structure of the RBDCRD-linker-NarX fusion proteins was predicted with AlphaFold2 from the amino acid sequence using the Latch Console platform (LatchBio). Using PyMOL (Version 2.5.4, Schrödinger, LLC.), we aligned the NMR-derived structure of two KRAS-RBDCRD dimers tethered to a nanodisc (worldwide Protein Data Bank, PDB accession code: 6PTS^71^) with the NMR structure of a KRAS4B-GTP homodimer on a lipid bilayer nanodisc (PDB accession code: 6W4E^72^). The predicted structure of two RBDCRD-linker-NarX proteins was then aligned to each of the RBDCRD structures. The flexible parts of the RBDCRD and the linkers were rotated, to bring the two NarX domains into proximity.

### Fluorescence Microscopy

Microscopy pictures were taken 36-48 h after transfection using a Nikon Eclipse Ti inverted microscope equipped with a Nikon Intensilight C-HGFI fiber illuminator, Semrock filter cubes (IDEX Health & Science), a 10x objective, and a Hamamatsu C10600 ORCA-R2 digital camera. Excitation filters, dichroic mirrors, emission filters, and exposure times are summarized in Supplementary Table 8. The Look-up table (LUT) values were adjusted for ideal contrast and kept constant within an experiment.

### Flow Cytometry

36-48h after transfection, we prepared the cells for flow cytometry analysis by removing the medium and adding 70-100 µL of Accuatase (Gibco, Thermo Fischer Scientific, cat.no. #A1110501). The cells were incubated for 15-30 min at room temperature and then kept on ice before measurement using a BD LSRFortessa^TM^ with a high-throughput screening device. To avoid potential cell damage and minimize time on ice, the plates were prepared consecutively, right before analysis. Excitation wavelengths, longpass filters, and emission bandpass filters were optimized to reduce crosstalk between different fluorophores and are summarized in Supplementary Table 9. When working with 24-well plates, after removing the medium, 150-200 µL of Accutase was used to detach the cells. After 15-30 minutes of incubation, the cells were detached by gentle pipetting, and the complete cell suspension was transferred to a 96-well plate for analysis using the HTS device.

### RAS-GTP Pulldown ELISA

RAS-GTP levels were measured using a Ras Activation ELISA assay kit (Merck Millipore cat.no. #17-497) according to the manual. HEK293 cells were seeded in 6-well plates and transfected as described in the ‘Transfection’ section. Cells were lysed using 250 µL of the provided lysis buffer with added Halt^TM^ Protease Inhibitor Cocktail (Thermo Fisher Scientific, cat.no. #89900). Samples were snap frozen in liquid nitrogen and stored at −80°C. An aliquot was used to quantify the protein amount in the cell lysates using a Pierce^TM^ BCA Protein Assay Kit (Thermo Fisher Scientific, cat.no #23227). The next day, 100 µg of each sample was used for the ELISA. The anti-RAS antibody (Merck Millipore cat.no. #17-497; part no.2006992) was provided in the ELISA kit. Chemiluminescence was measured using a Tecan Spark Multimode Microplate Reader and measured 20 min after the addition of the substrate with an integration time of 1 s.

### Confocal Microscopy for Quantification of Membrane Binding

Cells were seeded in a 24-well plate and transfected as described in the ‘Transfections’ section. After 36 h we detached the cells using 0.25% trypsin and re-seeded 30,000 cells in 200 µL medium into an 8-well glass-bottom plate to have sparsely distributed cells suitable for membrane detection and image analysis. The cells were incubated at 37°C, 5% CO2 to reattach. After 4 h, we stained the membrane of the cells with the CellBrite Steady 685 Membrane Staining Kit (20 µL of 1:100 dilution; Biotium cat.no. #30109-T) and the nuclei with 5 µL NucBlue Live Cell Stain (Thermo Fischer Scientific, cat.no. R37605). The wells were imaged using a Falcon SP8 confocal microscope (Leica Microsystems) with a 20x objective. SYFP2 was measured using an excitation laser at 524 nm and 540-600 nm bandpass filter, NucBlue was measured using the 405 nm laser and 415-460 nm bandpass filter, CellBrite was measured using a 670 nm excitation laser and a 750-800 nm bandpass filter.

### Image analysis for Quantification of Membrane Binding

Confocal images were analyzed with Python v3.10 using the Scikit-Image v0.19.2, SciPy v1.8.1, NumPy v1.22.4, Pandas v1.4.2, OpenCV v4.5.0 and custom packages built upon these libraries. Cells were segmented based on the membrane (CellBrite Steady 685) and nuclear (NucBlue) signals. The nuclear signals were used to assign the membrane signal to the individual cell. First, each nucleus was identified as a single cell. Second, the membrane signal was assigned to the nearest nucleus and used to create a mask for the membrane of each cell. Third, the membrane masks were post-processed to exclude non-closed objects, such as cell debris or cells with incomplete membrane staining as well as cells with more than one nucleus per cell. Fourth, the membrane masks were filled to additionally obtain the full cell masks. Fifth, for signal analysis, the total fluorescence in the SYFP2 channel was calculated for both the membrane and the full cell mask of each cell. Finally, the ratio between the membrane and the total cell signal was calculated for each cell. We used untransfected cells stained with NucBlue and CellBright steady 685 to assess background fluorescence. For the analysis, we included all transfected cells with total cell fluorescence above this background.

### Regression Model

The effect of different circuit parts on the output expression in HEK-293 with KRAS^G12D^, in HEK-293 with KRAS^WT^ as well as on the dynamic range (KRAS^G12D^/KRAS^WT^) was modeled using a Gaussian generalized linear model in R Studio v2023.06.0+421. Because the datasets were skewed toward low values, resulting in a loss of model accuracy at low values (Supplementary Fig. 15), a log transformation of all values was performed before running the models. Linearity and homoscedasticity assumptions were assessed using residual plots (Supplementary Fig. 16a-f). The goodness of fit was evaluated by comparing the fitted and measured values and calculating Pearson (R^2^) and Spearman correlations using the cor() function in R Studio (Supplementary Fig. 16g-i). For the selection of the independent variables, we tested three models assuming different interactions between the response elements and the parts they express. (Supplementary Table 10). The model with the best correlation (Model 3) was selected to analyze the effect of the different circuit parts in Figure 5c. In addition to the primary variables of interest, we considered potential covariates such as experimental batch, transfection efficiency, and amount of NarX and NarL plasmids. While the experimental batch did not affect the correlation of the model, transfection efficiency and plasmid amounts were included in the regression to ensure the robustness of the model.

### Analysis of Flow cytometry data

Flow cytometry data analysis was performed using FlowJo software v10.8.0 (BD Life Sciences). The gating strategy is shown in (Supplementary Fig. 17a-d). When multiple cell lines with different transfection efficiencies were used, the cells were binned using the expression of the mCherry transfection control to include only cells with high transfection levels > 10^3^ (Supplementary Fig. 17d-e). We further validated that there was no direct correlation between transfection efficiency and normalized output in our experimental data (Supplementary Fig. 18). Absolute units (AU) of fluorescence were calculated by multiplying the number of positive cells (frequency of parent) with the mean expression of the fluorophore:

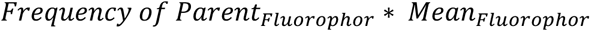

Relative units (RU) of fluorescence were calculated by dividing the AU of the fluorophore of interest by the AU of the transfection control:

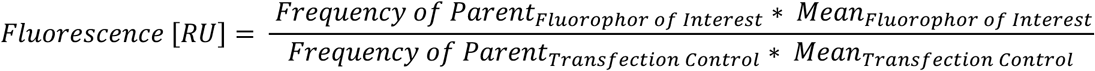

For the dynamic range, the fold change between the target cell line (HEK^G12D^ or HCT-116^WT^; ON-state) and off-target cell line (HEK^G12D^ or HCT-116^WT^; OFF-state) was calculated. The error bars were calculated using error propagation rules:

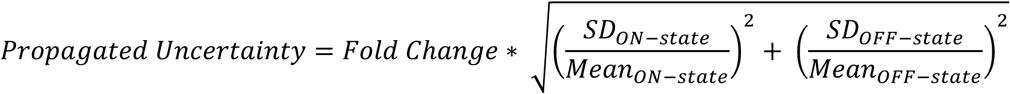

The flow cytometry histograms in Fig. 5d and Fig. 6e show all mCerulean-positive cells concatenated from the three biological replicates. The numerical values of the cell number, mean and frequency of parent can be found in Supplementary Tables 11 & 12.

### Analysis and Visualization of Screening Data

Data analysis and plotting of the data from the screening of different circuit parts (Fig. 5) and the screening of cancer cell lines (Fig. 7) were performed using R studio v2023.06.0+421. The data was imported from FlowJo analysis and labeled with the corresponding sample descriptors before calculating the mean and SD for the biological replicates. Plots were generated using ggplot2 v3.4.3 and cowplot v1.1.1. Large language models (ChatGPT3.5, ChatGPT4.0; OpenAI) were used to facilitate code writing.

### HSV-TK Killing Assays

Cells were transfected as described under transfection, except that only half the number of cells were seeded, to adjust for the long duration of the assay. 46h after seeding (ca. 36h after transfection), 100 µL of medium with GCV (Sigma, cat.no. SML2346-1ML) or was added to reach a final concentration 50 µM. Brightfield and mCherry confluences of the cells were continuously imaged over the time course of the assay using the xCelligence eSight Real-Time Cell Analysis (Agilent, USA, CA). The Agilent RTCA eSight software v.1.3.2 was used to create brightfield and red fluorescence segmentation masks and quantify the confluence. Segmentation parameters were selected for each cell line to optimize for cell size and background-to-cell contrast (Supplementary Table 13). Quantified confluence data was exported and analyzed using R Studio v2023.06.0+421. To adjust for differences in confluence before addition of GCV between the conditions, the confluence shown in Fig. 8 was background adjusted to the EF1a_HSV-TK condition of the same cell line at t=46h:

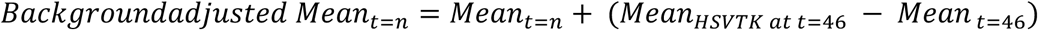

### Statistical Analysis

In experiments comparing two groups, unpaired two-tailed Student’s t-tests were used to assess significance (Fig. 1c & 7a and Supplementary Fig. 7d-I & 13a). When comparing three or more groups, an ordinary one-way ANOVA was used followed by a Dunnett’s multiple comparison test when all groups were compared with a control column (Fig. 2 & 8 and Supplementary Fig. 2b-f, 5 & 7a-c) or a Tukey’s test when all columns were compared (Supplementary Fig. 6). Significance in the regression model (Fig. 5c) was tested using a Wald test. Data was considered statistically significant at a p-value below 0.05. The number of replicates is provided in the figure captions. Each replicate was taken from a distinct sample. Apart from the large screening experiments (Fig. 5 & 6), data is representative of at least two experiments. The p-values, F-values, t-values, and degrees of freedom (df) from all statistical comparisons are provided in Supplementary Table 14. Statistical analyses were performed using Prism 10 (GraphPad) or R Studio (v2023.06.0+421).

## Data and Code availability

All data supporting the findings and the code used to calculate membrane localization, analyze the screenings, and for the regression model are available from the corresponding author upon request.

## Acknowledgments

This work was supported by the National Center of Competence in Research Molecular Systems Engineering (NCCR-MSE) and ETH Zurich core funding. We thank M. Di Tacchio, C. Cavallini, A. Gumienny, E. Montani, and A. Ponti for assistance with flow cytometry and image acquisition as well as analysis. We also thank B. Treutlein for discussions, proofreading, and providing the lab infrastructure during the last part of the project. We thank D. Schweingruber, F. Trick, V. Cheras, and S. Seidel for proofreading. Lastly, we would like to thank A. Abraham, J. Jaekel, J. Schreiber, M. Dastor, M. Lampis, P. Müller-Thümen, and all Benenson and Treutlein lab members for discussions.

## Author Contributions

G.S. conceived the study, performed the majority of experiments, analyzed the data, and wrote the manuscript. L.N. planned and performed some experiments and analyzed data. Y.B. conceived the study, analyzed data, wrote the manuscript, and supervised the project.

## Competing Interests

YB is a shareholder and an employee of Pattern Biosciences

## Supplementary Figures

**Supplementary Figure 1.**
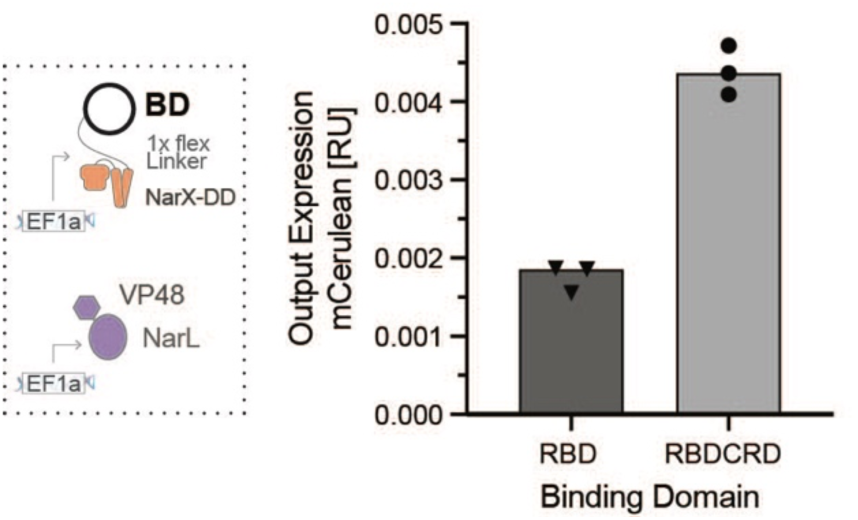
RAS Sensor activation with RBD or RBDCRD as binding domain. mCerulean output expression of the RAS Sensor with either RBD or RBDCRD of CRAF as binding domain in the NarX fusion proteins in HEK293 cells co-transfected with 15 ng of KRAS^G12D^. mCerulean expression was measured using flow cytometry and normalized to a constitutively expressed mCherry transfection control. Mean values were calculated from three biological replicates.

**Supplementary Figure 2.**
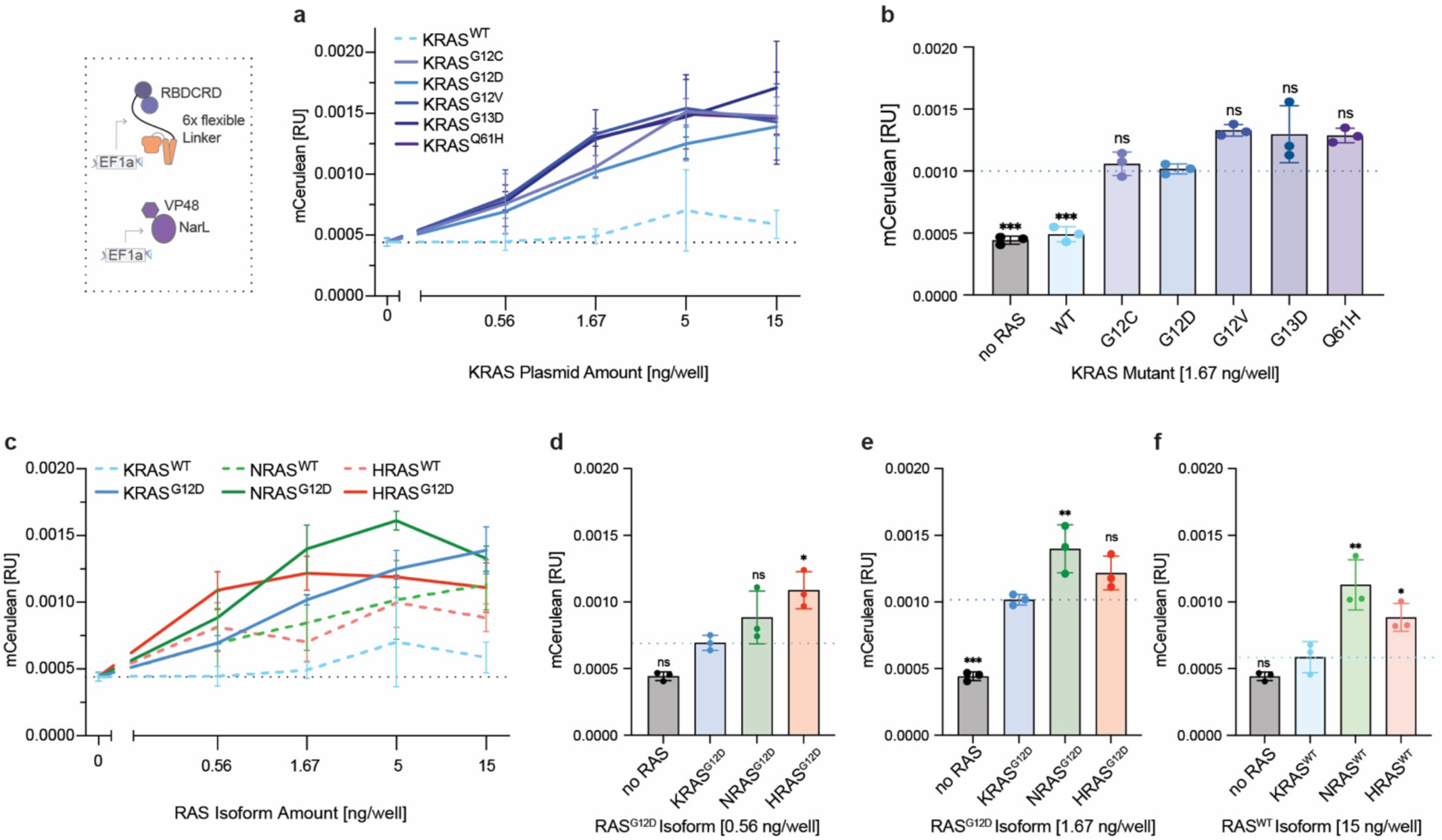
RAS Isoforms and Mutants. (**a**) Comparison of oncogenic KRAS mutants. Output expression of the RAS sensor when co-transfecting increasing amounts of different KRAS mutants. The dashed line represents wildtype KRAS (**b**) Bar chart of output expression at 1.67 ng KRAS plasmid/well from a. (**c**) Comparison of RAS isoforms. Output expression of the RAS sensor when co-transfecting increasing amounts of KRAS (blue), NRAS (green) or HRAS (red). The solid line represents G12D mutant, dashed line the corresponding wildtype isoform (**d**) Bar chart of output expression at 0.56 ng plasmid/well of different RAS^G12D^ Isoform from c. (**e**) Bar chart of output expression at 1.67 ng plasmid/well of different RAS^G12D^ Isoform from c. (**f**) Bar chart of output expression at 15 ng plasmid/well of different wildtype RAS Isoform from c. mCerulean output expression was measured by flow cytometry and normalized to a constitutively expressed mCherry transfection control. Mean values were calculated from biological triplicates. Error bars represent +/- SD. Significance was tested using an ordinary one-way ANOVA with Dunnett’s multiple comparisons to compare each condition with KRAS^G12D^ (b, d & e) or KRAS^WT^ (f). ns = non-significant, *p < 0.05, **p < 0.01, ***p < 0.001, ****p < 0.0001.

**Supplementary Figure 3.**
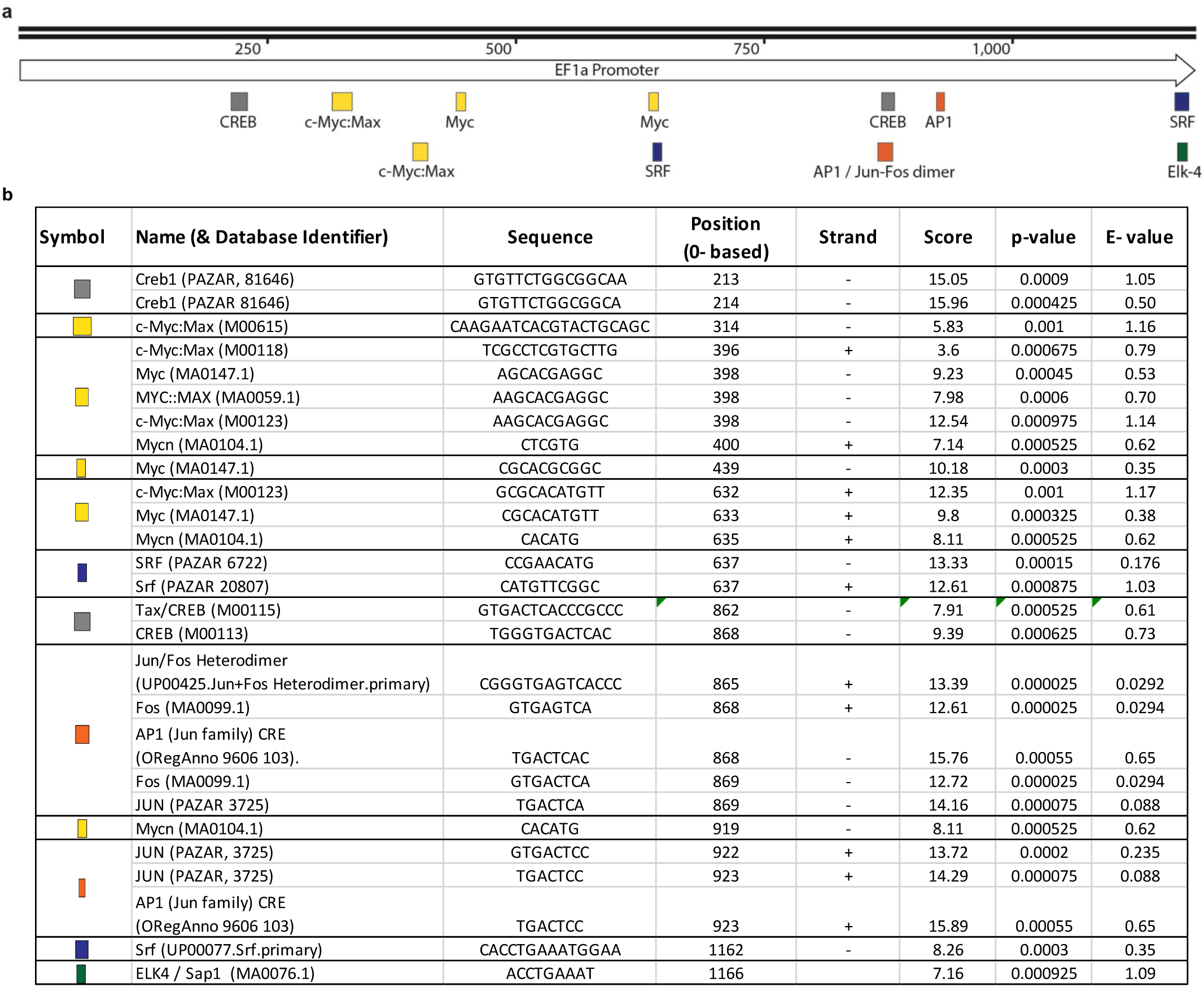
Prediction of MAPK transcription factor binding sites in EF1a. (**a**) Elongation factor 1a promoter (EF1a) with localization of predicted binding sites of MAPK transcription factors downstream of RAS. The width of the symbols indicates the sequence length of the potential binding site. (**b**) List of all predicted binding sites of Myc, AP1 (Fos/Jun), SRF, CREB, and Ets/Elk transcription factors with LASAGNA 2.0 using the TRASFAC TFBS search. For each binding site the sequence, position in EF1a, strand, binding strength (score), statistical significance (p-value), and expected number of random hits with similar scores in the data set (e-value) are provided.

**Supplementary Figure 4.**
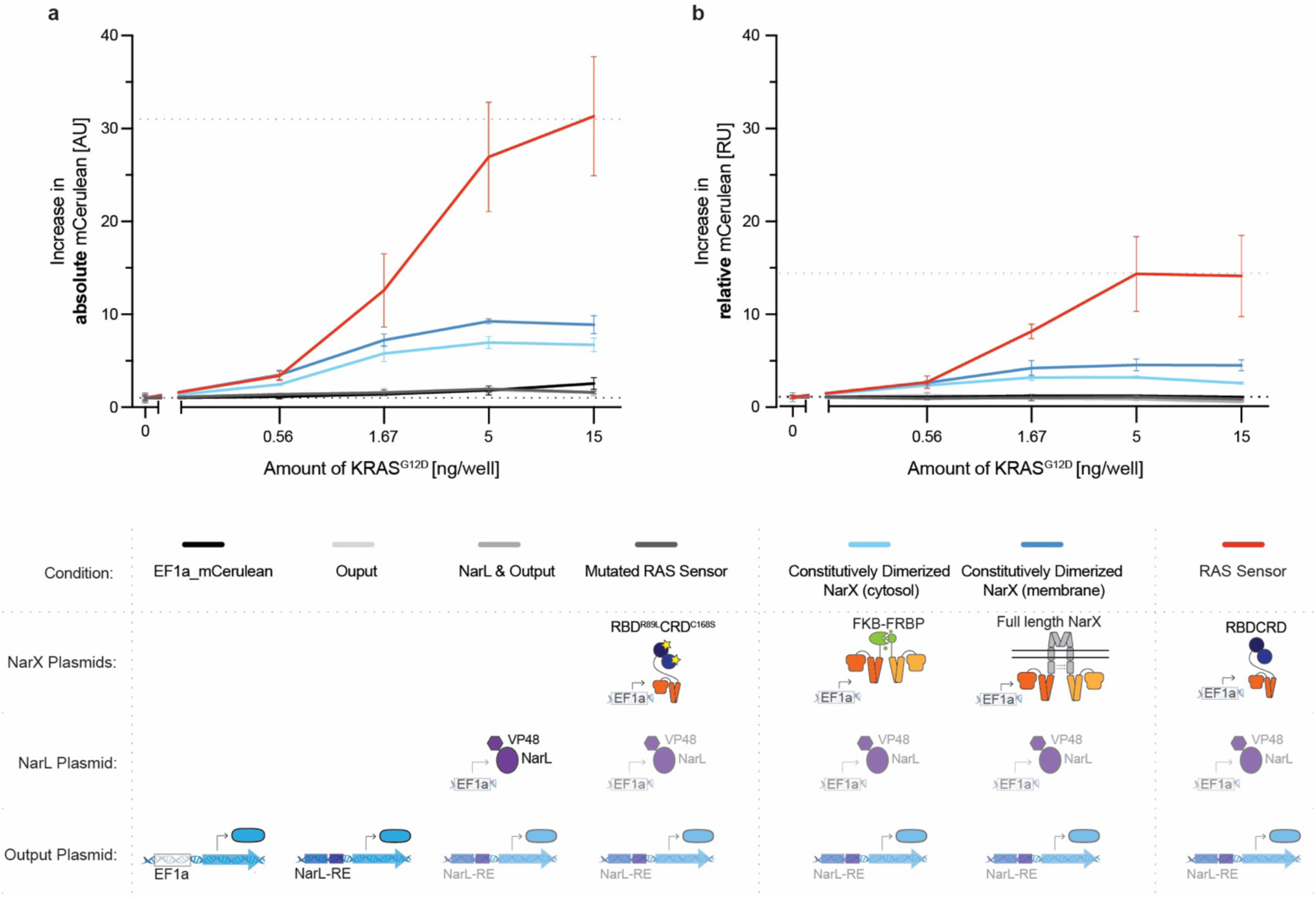
Contribution of RAS-dependent Increase in Sensor Expression Levels on total Output. Comparison of the functional RAS sensor (red) with inactive non-functional (different shades of gray) and constitutively active sensor controls (shades of blue). The details of the controls are illustrated in the table below. Black: constitutive EF1a driven mCerulean expression; light gray: only output plasmid present; gray: EF1a-driven NarL-VP48 expression and output plasmid present; dark gray: EF1a-driven expression of RAS-binding impaired mutated RBD^R89L^CRD^C168S^-NarX fusion proteins and NarL-VP48 and output plasmid present; light blue: same plasmids as RAS sensor but RBDCRD in the NarX fusion proteins are replaced by FKB or FRBP, which constitutively dimerized in the cytosol (100 nM of A/C heterodimerized were added to induce full dimerization); dark blue: same plasmids as RAS sensor but the NarX fusion proteins were replaced by a full length NarX protein that constitutively dimerizes at the membrane. (**a**) Increase in absolute mCerulean signal (non-normalized) compared to 0 ng/well of KRAS^G12D^. (**b**) Increase in relative mCerulean signal (= mCerulean output expression normalized to a constitutively expressed mCherry) compared to 0 ng/well of KRAS^G12D^ plasmid. Mean values were calculated from biological triplicates. Error bars represent +/- SD.

**Supplementary Figure 5.**
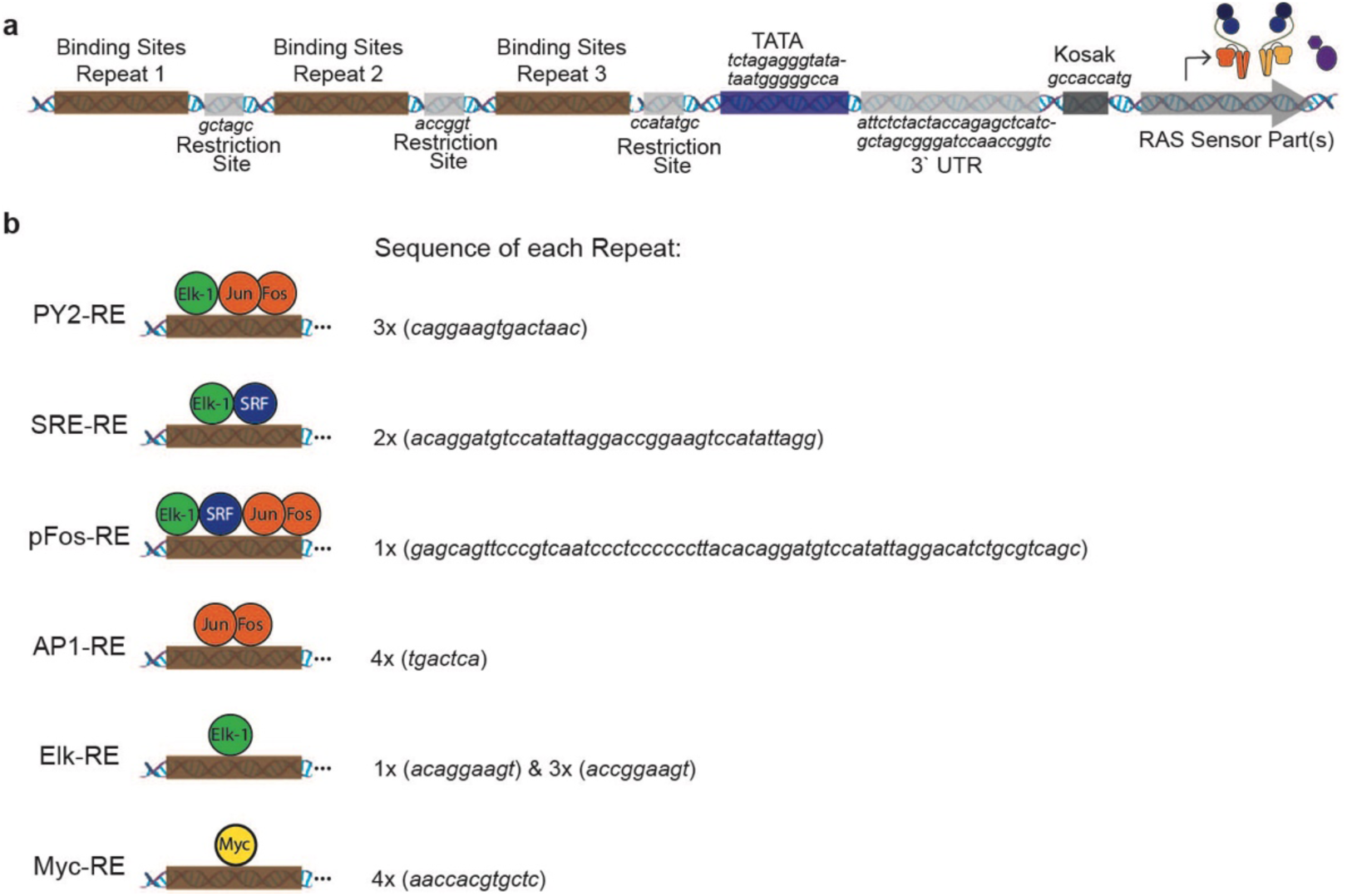
Design of the MAPK Response Elements. (**a**) Detailed representation of the MAPK response elements design. Three repeats of the transcription factor binding site pattern with restriction enzyme cutting sites were placed in front of a minimal promoter (TATA). Exact sequences of the restriction sites, TATA, 3’ UTR, and Kosak are shown in italics. (**b**) Sequences of the transcription factor binding site of each repeat of the different response elements. The schematics show which transcription factors bind to which response element.

**Supplementary Figure 6.**
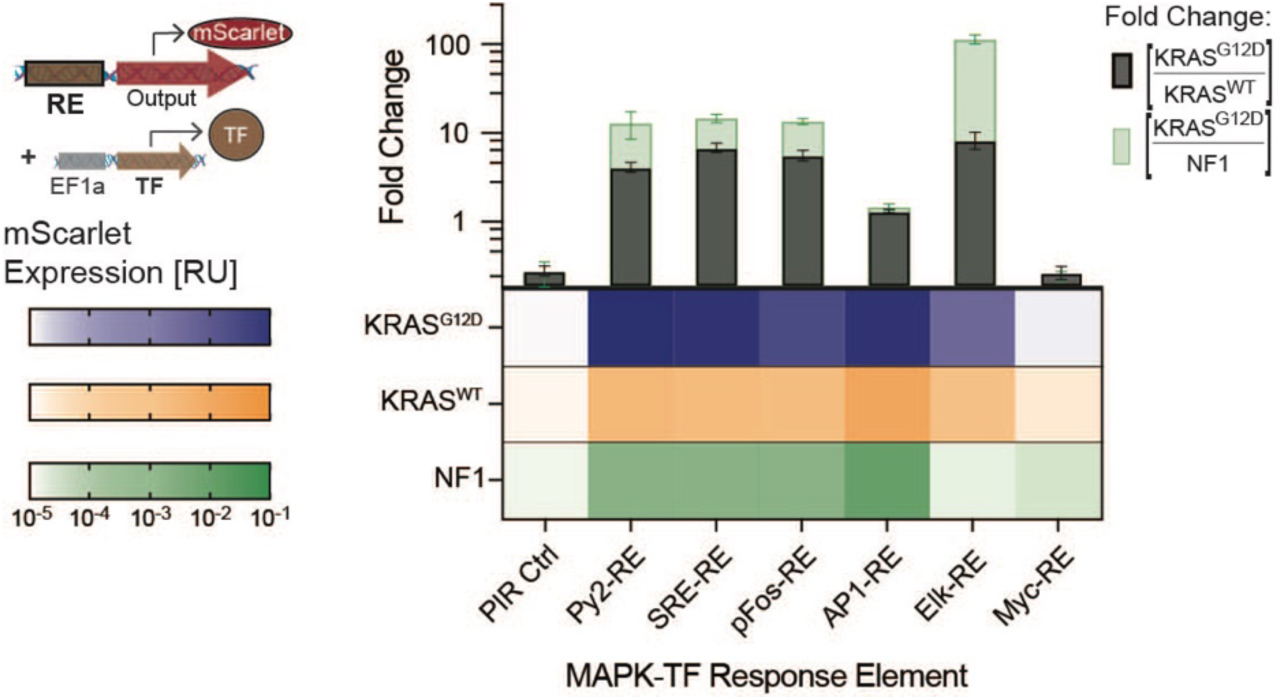
MAPK Response Element with overexpressed transcription factors. mScarlet expression from plasmids with the different MAPK response elements when additionally overexpressing the MAPK transcription factors Elk-1, c-Jun, c-Fos, and c-Myc. Heatmap showing the normalized mScarlet expression in HEK293 cells co-transfected with 15 ng/well of either KRAS^G12D^ (blue), KRAS^WT^ (orange), or NF1, a GTPase-activating protein that deactivates endogenous RAS (green). Bars above show the corresponding fold changes between cells with KRAS^G12D^ and KRAS^WT^ in black or KRAS^G12D^ and NF1 in green. mCerulean expression was measured by flow cytometry and normalized to a constitutively expressed mCherry transfection control. Mean values were calculated from two biological replicates.

**Supplementary Figure 7.**
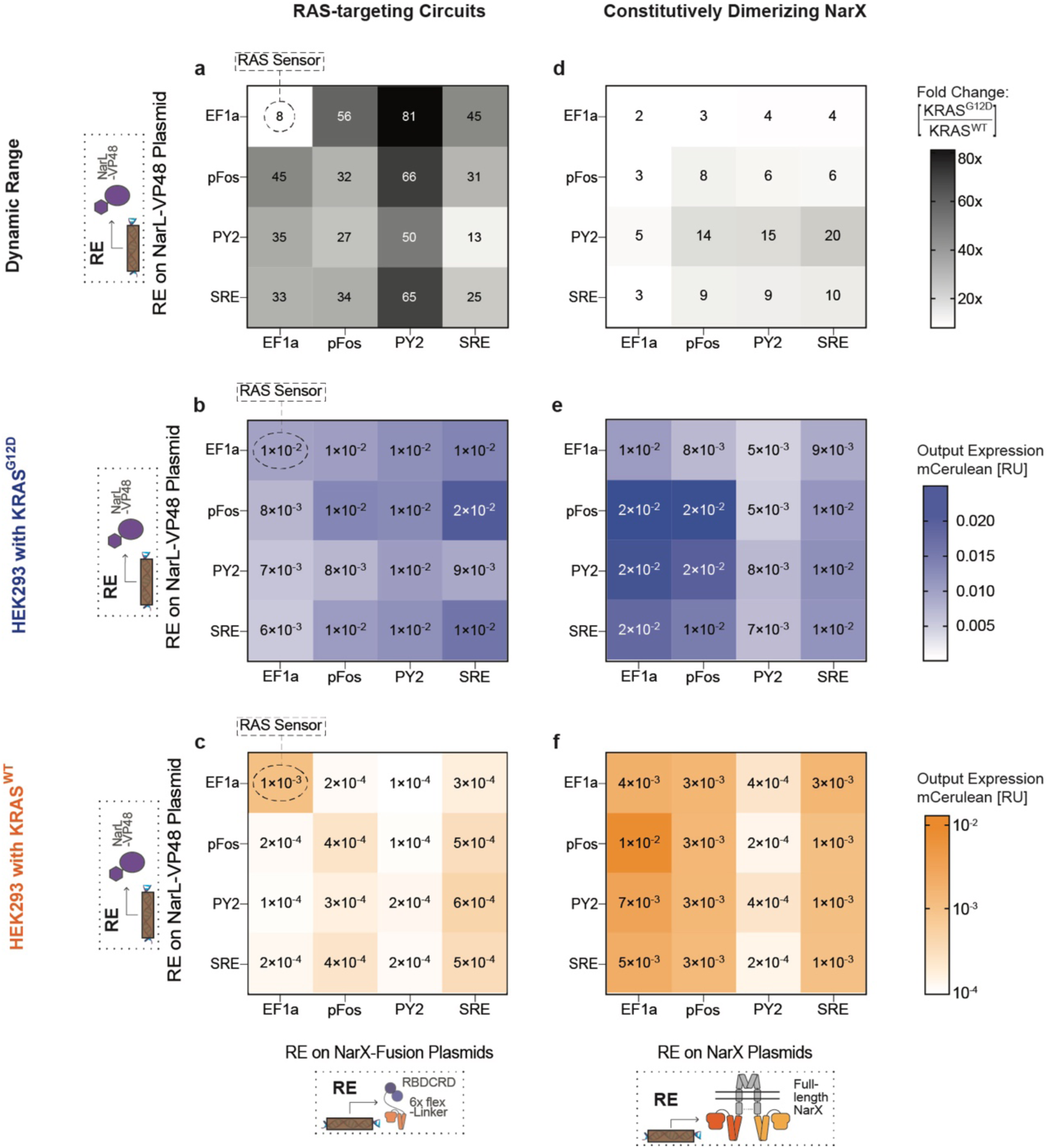
Comparison of the RAS-binding dependent AND-gate RAS circuits and MAPK response element expressed constitutively dimerized NarX-NarL TCS. (**a**) Dynamic range of the RAS-targeting circuits. Fold change in mCerulean output expression between HEK293 co-transfected with 1.67 ng/well of KRAS^G12D^ and KRAS^WT^. In the RAS targeting circuits, NarL-VP48 and/or the RBDCRD-6xfL-NarX fusion proteins were expressed using different MAPK-REs or a constitutive promoter (EF1a). (**b)** Output expression of RAS-targeting circuits with different response elements in HEK293 co-transfected with 1.67 ng/well of KRAS^G12D^. (**c**) Output expression of RAS-targeting circuits with different response elements in HEK293 co-transfected with 1.67 ng/well of KRAS^WT^. (**d-f)** Same set-up as in a-c, but the RBDCRD-NarX fusion protein sequence on the NarX plasmids were replaced by full length NarX that constitutively dimerizes at the membrane. Fluorescent protein expression was measured by flow cytometry and normalized to a constitutively expressed transfection control. Mean values were calculated three biological replicates. PY2: polyoma virus enhancer domain; SRE: Serum response element; pFos: minimal promoter of c-fos; AP1: activator protein 1; Elk: Ets-like protein; Myc: myelocytomatosis protein.

**Supplementary Figure 8.**
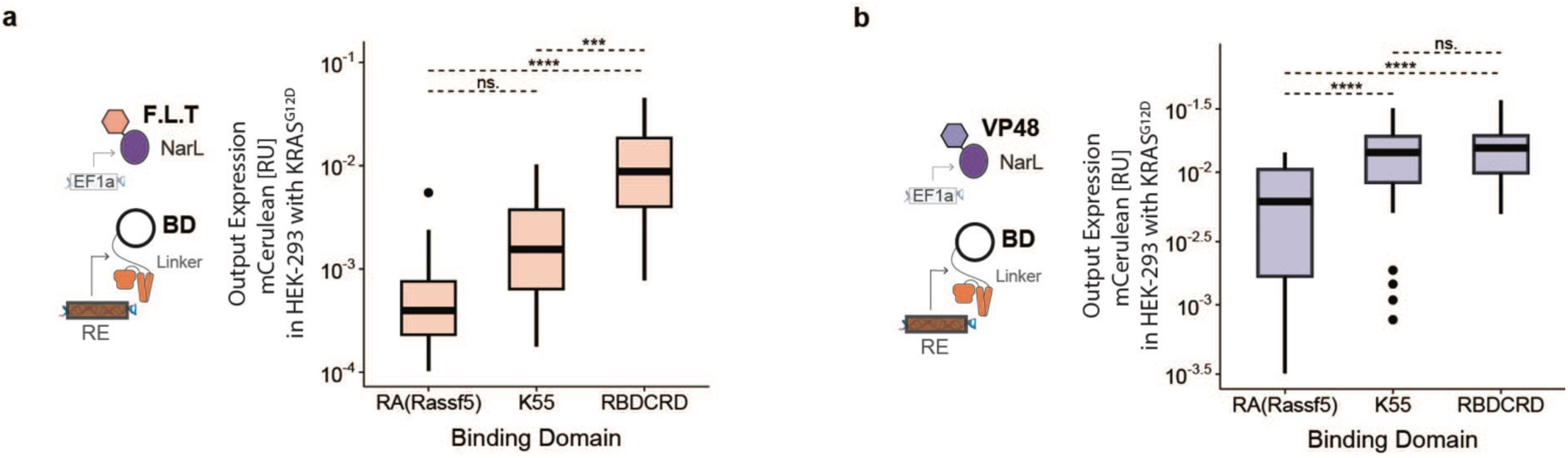
Differential circuit activation of the binding domains (BD) depends on the transactivation domain fused to NarL. Boxplots of mCerulean output expression in HEK293 co-transfected with 1.67 ng of KRAS^G12D^. The boxplots include all RAS-targeting circuits from the screening in Fig.5, where the NarX fusion proteins were expressed through the response elements split by the binding domain used in the circuit. (**a**) Effect of binding domain in circuits with F.L.T. as transactivation domain fused to NarL. (**b**) Effect of the binding domain in circuits with VP48 as transactivation domain. mCerulean expression was measured by flow cytometry and normalized to a constitutively expressed mCherry transfection control. Each of the boxplot shows all circuits from the screening that used the indicated binding and transactivation domain, resulting in a total of 18 circuits in **a** or 34 in **b**. For each circuit three biological replicates were measured. Significance was tested by one-way ANOVA with a Tukey’s multiple comparison. *p < 0.5, ***p < 0.001, ****p < 0.0001.

**Supplementary Figure 9.**
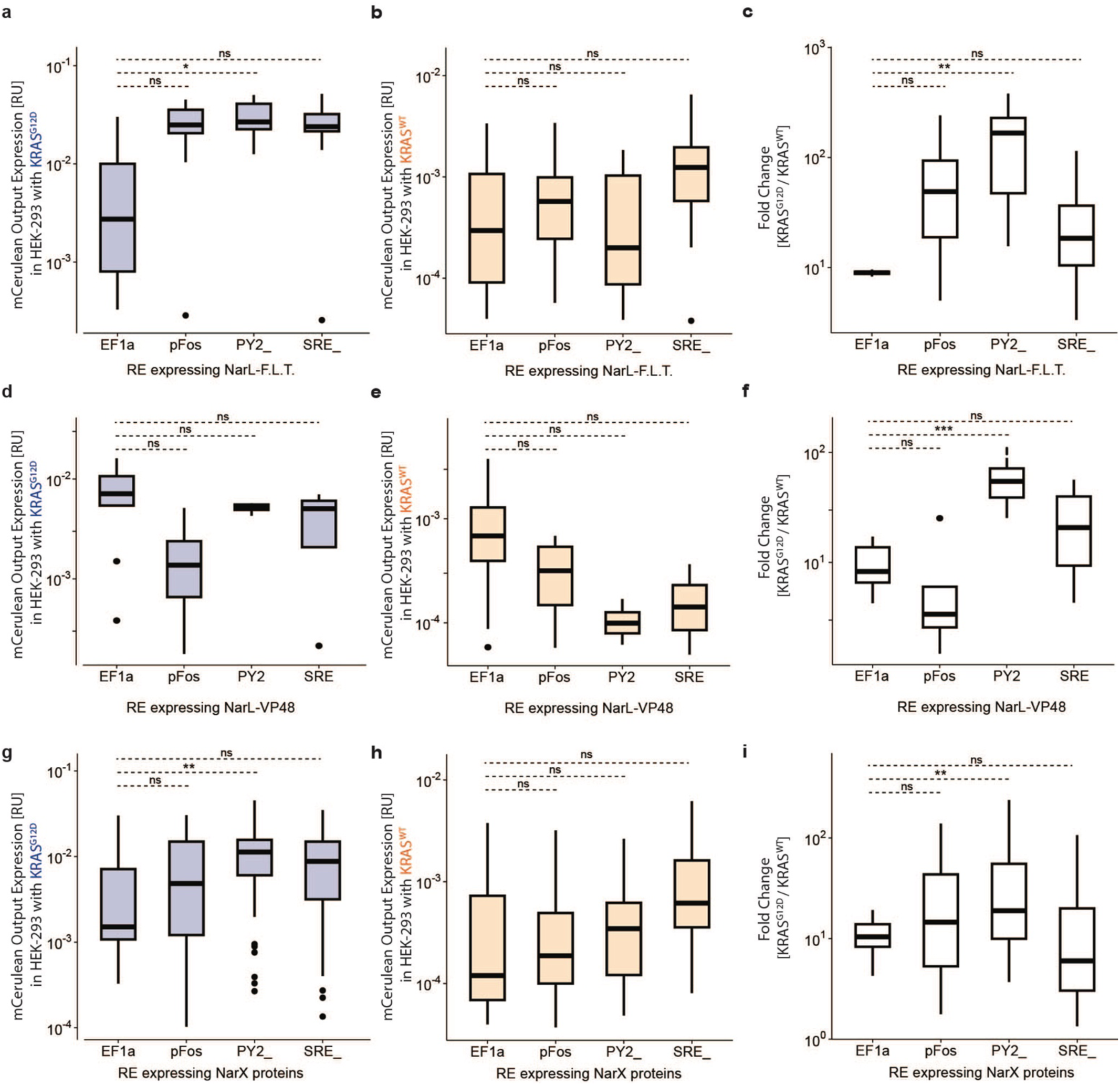
Effect of the RAS circuit parts: response elements (RE). Boxplots of activation of all RAS-targeting circuits with either EF1a as promoter or pFos, Py2, or SRE as response element expressing different circuit parts. (**a-c**) Effect of response elements when expressing NarL-F.L.T. via REs, while the NarX fusion proteins are expressed with EF1a. (**d-f**) Effect of response elements when expressing NarL-VP48 via REs, while the NarX fusion proteins are expressed with EF1a. (**g-i**) Effect of response elements when expressing NarX fusion proteins via REs, while NarL is expressed with EF1a. **a,d&g** show the mCerulean output expression in HEK293 co-transfected with 1.67 ng of KRAS^G12D^ plasmid. **b,e&h** show the mCerulean output expression in HEK293 co-transfected with 1.67 ng of KRAS^WT^ plasmid. c,f&i show the fold change between cells with KRAS^G12D^ and KRAS^WT^. mCerulean expression was measured using flow cytometry and normalized to a constitutively expressed mCherry transfection control. Mean values were calculated from three biological replicates. Each of the Boxplot shows all circuits from the screening that used the indicated response element to express the indicated part, resulting in a total of 4 circuits in **a-c**, 14 in **d-f,** or 48 in **g-i**. For each circuit three biological replicates were measured. Significance was tested using a one-way ANOVA with a Dunnet’s multiple comparison against the EF1a condition. *p < 0.5, **p < 0.01, ***p < 0.001.

**Supplementary Figure 10.**
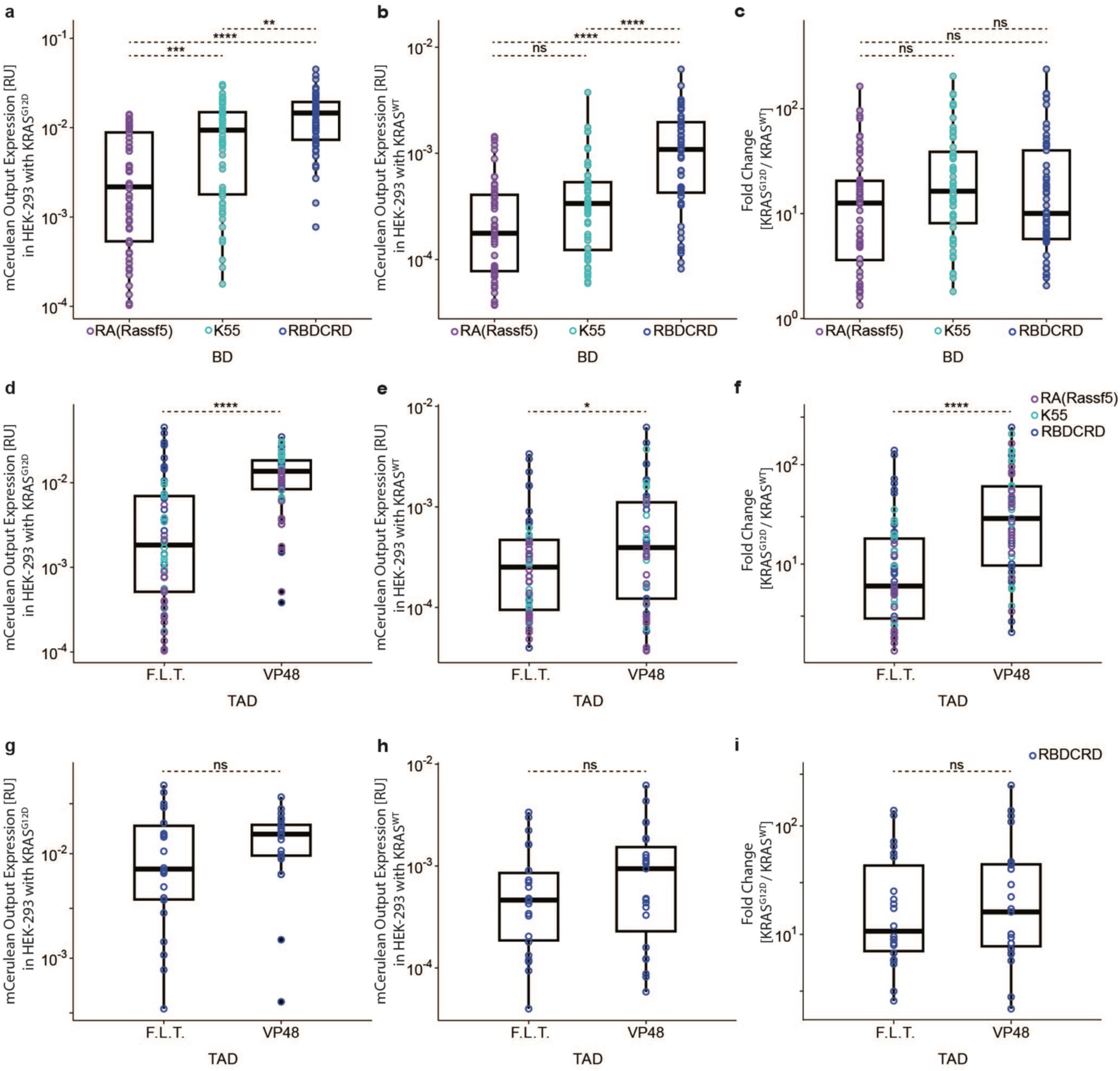
Effect of the RAS circuit parts: binding domain (BD) and transactivation domain (TAD) in RE_NarX circuits. (**a-c**) Effect of the binding domain. Boxplots show the mCerulean output expression in HEK293 of all tested RAS-targeting circuits where the NarX fusion proteins are expressed via MAPK response elements (RE_NarX circuits) split by the used binding domain. **a** shows the ON-state in cells co-transfected with KRAS^G12D^, **b** the OFF-state in cells co-transfected with KRAS^WT,^ and **c** the dynamic range calculated as fold change between ON- and OFF-state. (**d-f**) Effect of the transactivation domain. Boxplots show mCerulean output expression of the same circuits as in **a-c** but split by the transactivation domain fused to NarL. **d** shows the ON-state, **b** the OFF-state, and **f** the dynamic range. (**g-i**) Effect of the transactivation domain only in circuits with RBDCRD as binding domain. Boxplots show mCerulean output expression in the circuits from **d-f** that use RBDCRD as binding domain. **g** shows the ON-state, **h** the OFF-state, and **i** the dynamic range. Each circle represents one circuit condition using one of the following binding domains: RA(Rassf5) (purple), K55 (cyan), or RBDCRD (blue). For each circuit three biological replicates were measured. mCerulean output expression was measured using flow cytometry and normalized to a constitutively expressed mCherry transfection control. Significance was tested using a one-way ANOVA with a Tukey’s multiple comparisons in **a-c** and unpaired two-tailed Student’s t-tests in **d-i**. *p < 0.5, **p < 0.01, ***p < 0.001.

**Supplementary Figure 11.**
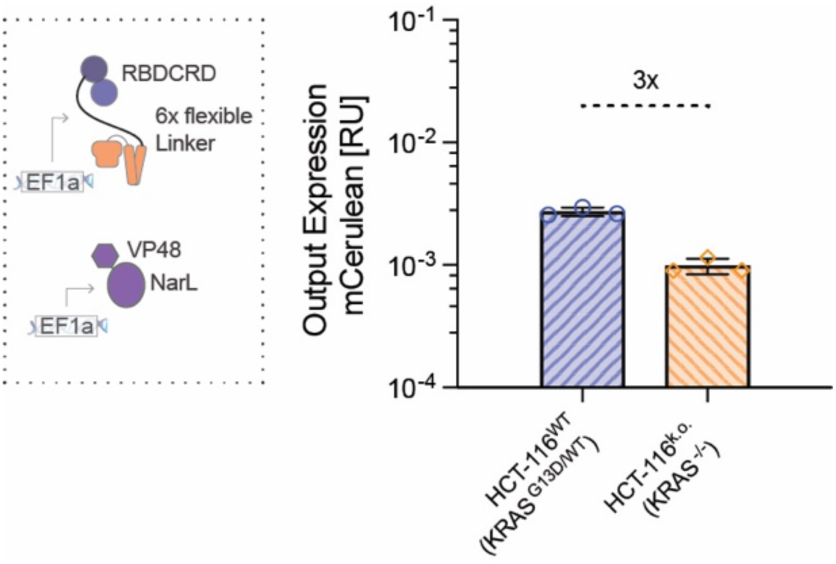
RAS Sensor activation in HCT-116. mCerulean output expression of the RAS Sensor in HCT-116 wildtype (blue) or KRAS knock out (orange) cells. mCerulean expression was measured using flow cytometry and normalized to a constitutively expressed mCherry transfection control. Mean values were calculated from three biological replicates.

**Supplementary Figure 12.**
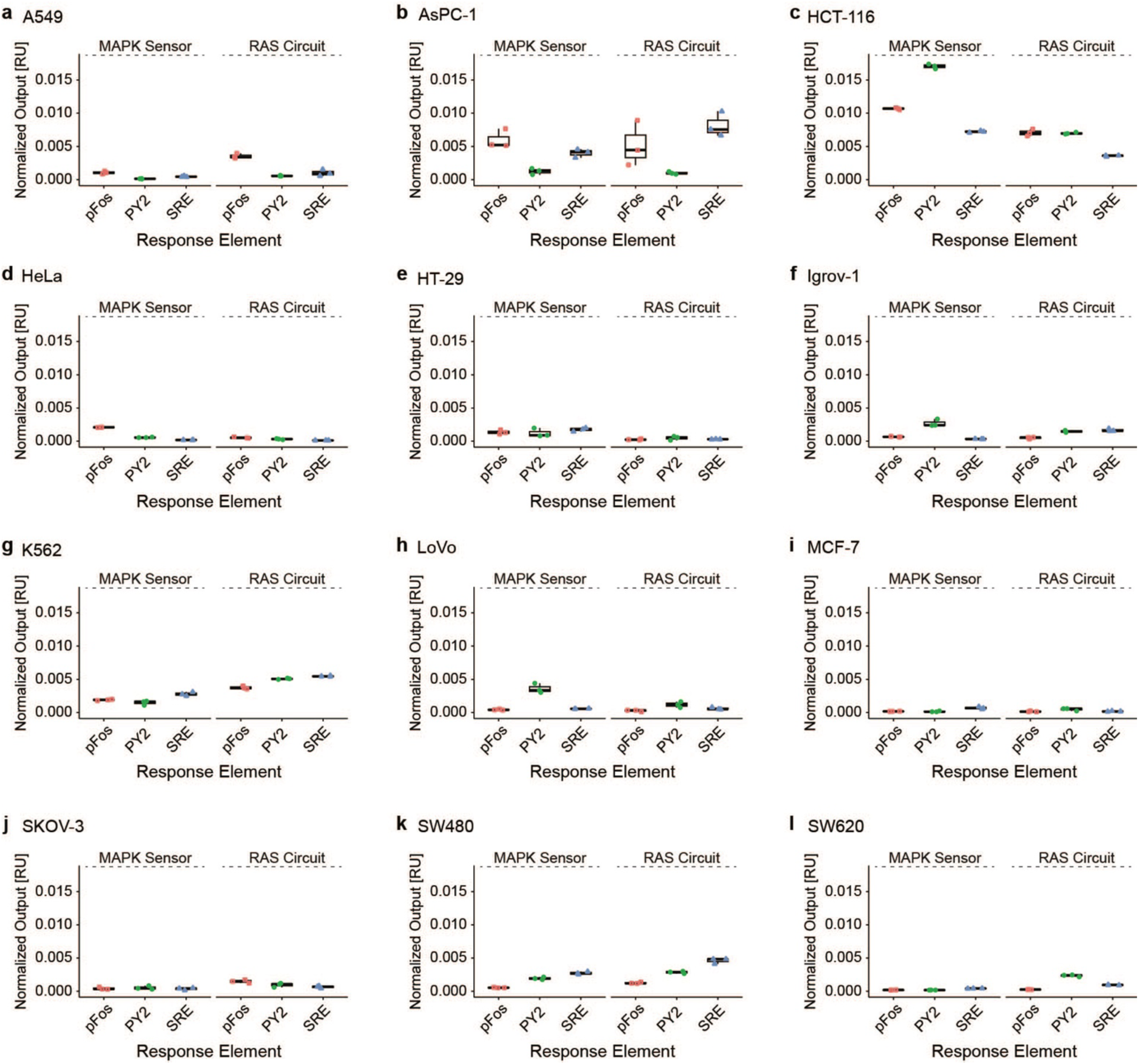
Correlation of activation of the MAPK sensors and the RAS-targeting circuits in individual cell lines. Output expression of the MAPK sensors or the RAS circuits when using pFos (red) PY2 (green) or SRE (blue) as response elements. For the MAPK sensors, the response elements directly expressed mScarlet. For the RAS Circuits, the response elements expressed NarL-F.L.T. leading to mCerulean output expression. **a** A549, **b** AsPC-1, **c** HCT-116, **d** HeLa, **e** HT-29, **f** Igrov-1, **g** K562, **h** LoVo, **i** MCF-7, **j** SKOV-3, **k** SW480, **l** SW620. Output expression was measured using flow cytometry and normalized to a constitutively expressed transfection control. Each symbol represents one biological replicate (n=3).

**Supplementary Figure 13.**
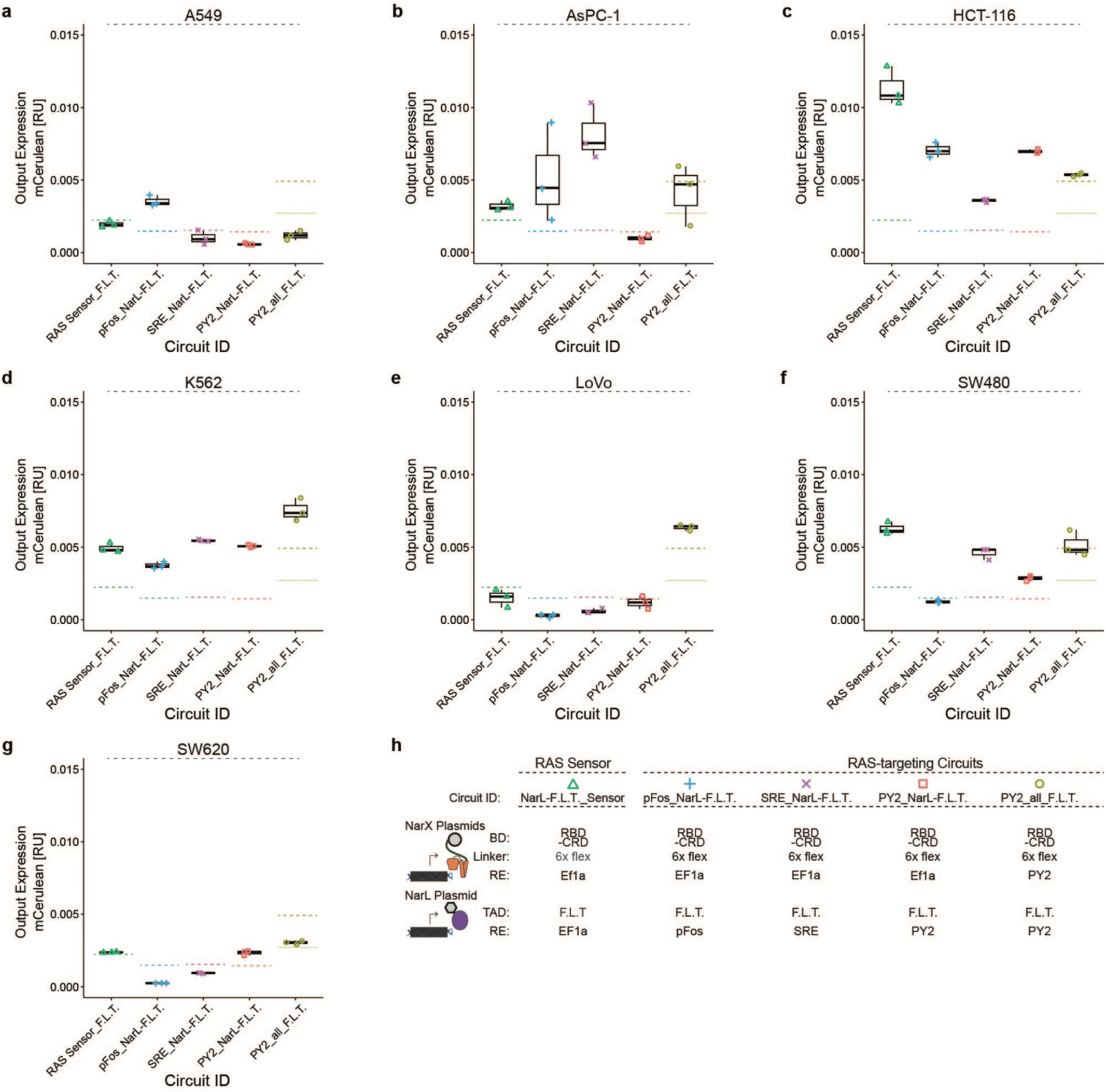
Comparison of RAS-targeting circuits in individual cancer cell lines with mutations overactivating RAS. Output expression after transfecting different RAS circuits into various cancer cell lines with mutations overactivating RAS. The dashed lines show the median activation of the RAS^WT^ cell lines with the highest background for the corresponding circuit, namely Igrov-1 for RAS Sensor_F.L.T. (green), SRE_NarL-F.L.T. (purple) and PY2_NarL-F.L.T. (red), SKOV-3 for pFos_NarL-F.L.T. (blue) and HT-29 for PY2_all_F.L.T. (olive). As HT-29 has a BRAF mutation overactivating the RAS pathway, for PY2_all_F.L.T. HeLa, the highest RAS^WT^ cell line without MAPK mutation, is also shown as a dotted line (olive). (**a**) A549, (**b**) AsPC-1, (c) HCT-116, (**d**) K562, (**e**) LoVo, (**f**) SW480, (**g**) SW620 cells. The composition of the parts used in the different RAS circuits is shown in **h**. mCerulean output expression was measured using flow cytometry and normalized to a constitutively expressed mCherry transfection control. Mean values were calculated from three biological replicates.

**Supplementary Figure 14.**
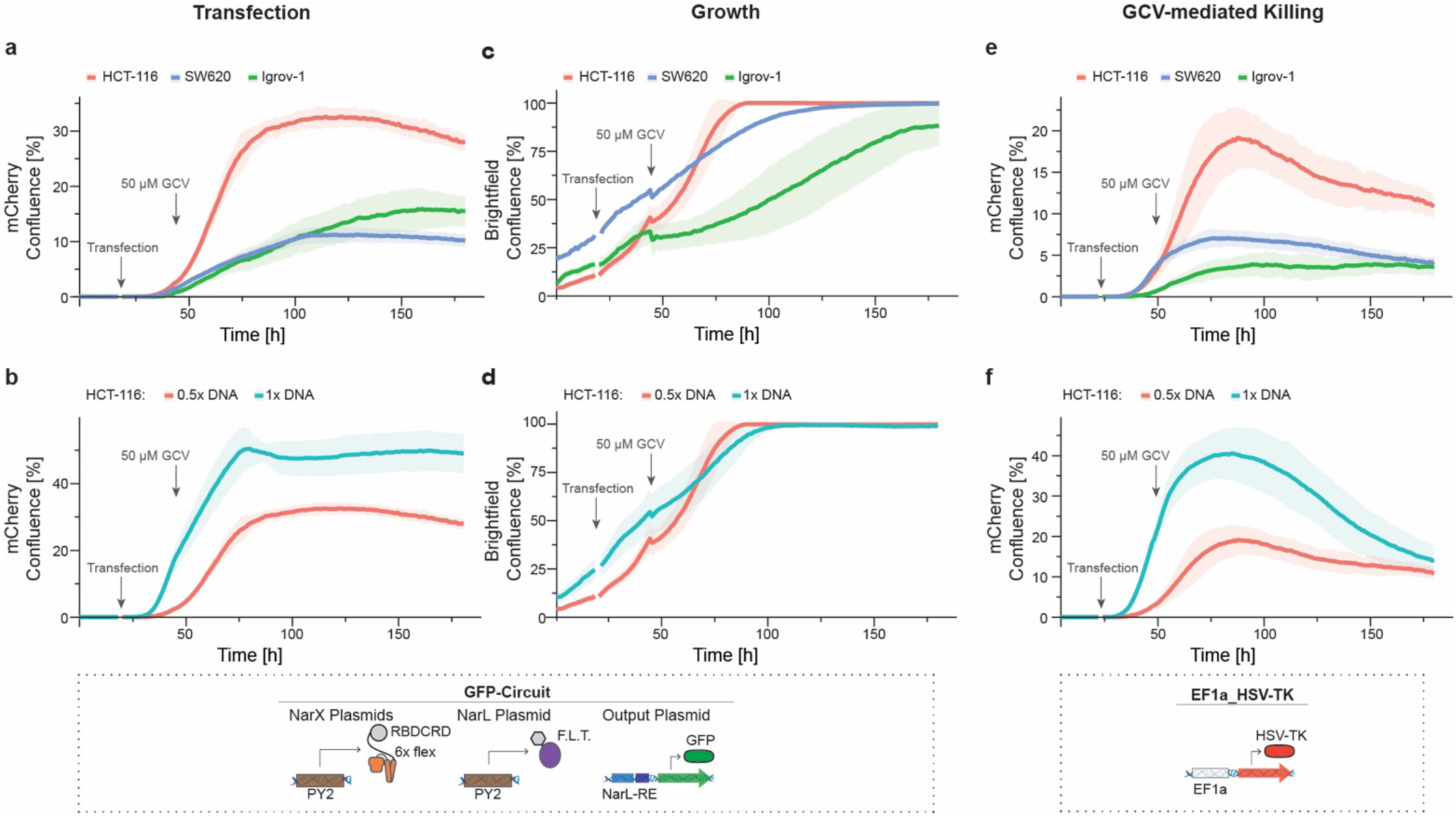
Differences between cell lines and transfection amount in killing assay. The graphs show the confluence of mCherry transfection control positive cancer cells (**a,b,e,f**) or overall confluence (**c,d**) transfected with a RAS circuit expressing GFP as output as negative control without HSV-TK (**a-d**) or a EF1a-expressed constitutive HSV-TK as positive control (**e-f**). Control circuits are illustrated in the dotted boxes below the graphs. Gray arrows indicate the timepoint of transfection and the addition of the prodrug ganciclovir (GCV) that is activated by HSV-TK. **a** Difference in transfection efficiency between cell lines measured as confluence of the constitutively expressed mCherry transfection control. **b** Difference in transfection efficiency in HCT-116 when transfecting different amounts of circuit DNA. **c** Different growth curves between cell lines. Growth was measured as overall confluence in the well; red = HCT-116 (0.5x DNA amount), blue = SW630, green = Igrov-1. **d** Growth curves in HCT-116 when transfecting different amounts of circuit DNA. Red = 0.5x DNA amount, Turquoise = 1x DNA amount. **e** Difference in killing curves between cancer cell lines when constitutively expressing herpes simplex virus thymidine kinase (HSV-TK). Killing of transfected cells is approximated using the confluence of mCherry positive cells. **f** Difference in killing curves in HCT-116 when transfecting different amounts of DNA. Confluence was quantified from microscopy images. Mean confluence was calculated from biological triplicates with standard deviation shown as ribbons. Significance was tested using an ordinary one-way ANOVA with Dunnett’s multiple comparison test. ns = non-significant, *p < 0.05, **p < 0.01, ***p < 0.001, ****p < 0.0001.

**Supplementary Figure 15.**
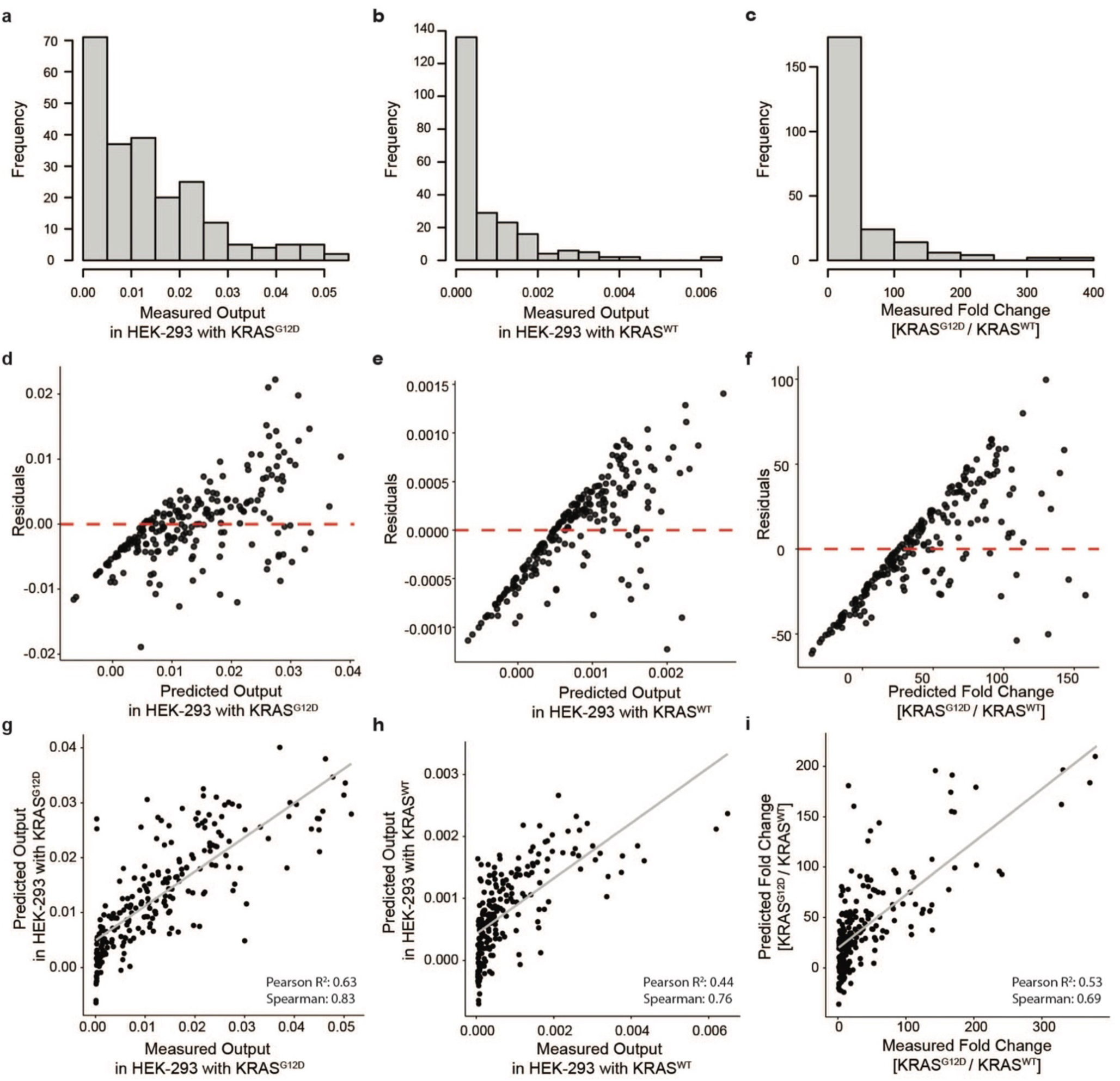
Assessment of the non-transformed regression model. (**a-c**) Histograms showing the distribution of the measured mCerulean output expression of all circuits tested in the screening in HEK293 overexpressing KRAS^G12D^ in **a**, KRAS^WT^ in **b**, or the fold change between KRAS^G12D^ and KRAS^WT^ in **c**. (**d-f**) Residuals plots showing the difference between measured and fitted output (residuals) vs the predicted output of HEK293 overexpressing KRAS^G12D^ in **d**, KRAS^WT^ in **e**, or the fold change between KRAS^G12D^ and KRAS^WT^ in **f** used to assess linearity and homoscedasticity assumptions. (**g-i**) Goodness of fit: Measurements of output vs predictions of the regression models based on the parts used in the circuit. Predicted values were calculated using the following general linear regression models. **a,d,g** show the ON-state data in HEK293 co-transfected with KRAS^G12D^. Model formula: RASmut_RU ∼ RASmut_mCherry_AU + BD + RE_NarL_TAD + Linker + RE_NarX + Conc_NarX + Conc_NarL. **b,e,h** show the OFF-state in HEK293 co-transfected with KRAS^WT^. Model formula: RASwt_RU ∼ RASwt_mCherry_AU + BD + RE_NarL_TAD + Linker + RE_NarX + Conc_NarX + Conc_NarL. **c,f,i** show the dynamic range calculated as fold change between ON- and OFF-state. Model formula: Ratio ∼ RASmut_mCherry_AU + RASwt_mCherry_AU + BD + Linker + RE_NarX + RE_NarL + Conc_NarX + Conc_NarL Gray lines in g-I display the linear best fit used for calculating the Pearson correlation. RASmut_RU: mCerulean signal with KRAS^G12D^ normalized to transfection control, RASwt_RU: mCerulean signal with KRAS^WT^ normalized to transfection control, Ratio: fold change between RASmut_RU & RAS_wt_RU, RASmut_mCherry_AU: absolute transfection signal with KRAS^G12D^, RASmut_mCherry_AU: absolute transfection signal with KRAS^WT^, BD: binding domain, Linker: linker used in NarX fusion protein, RE_NarL_TAD: response element used to express NarL and transactivation domain fused to NarL, RE_NarX: response element used to express NarX proteins. Conc_NarX: transfected amount of plasmid expressing the NarX proteins. Conc_NarL: transfected amount of plasmid expressing the of NarL protein.

**Supplementary Figure 16.**
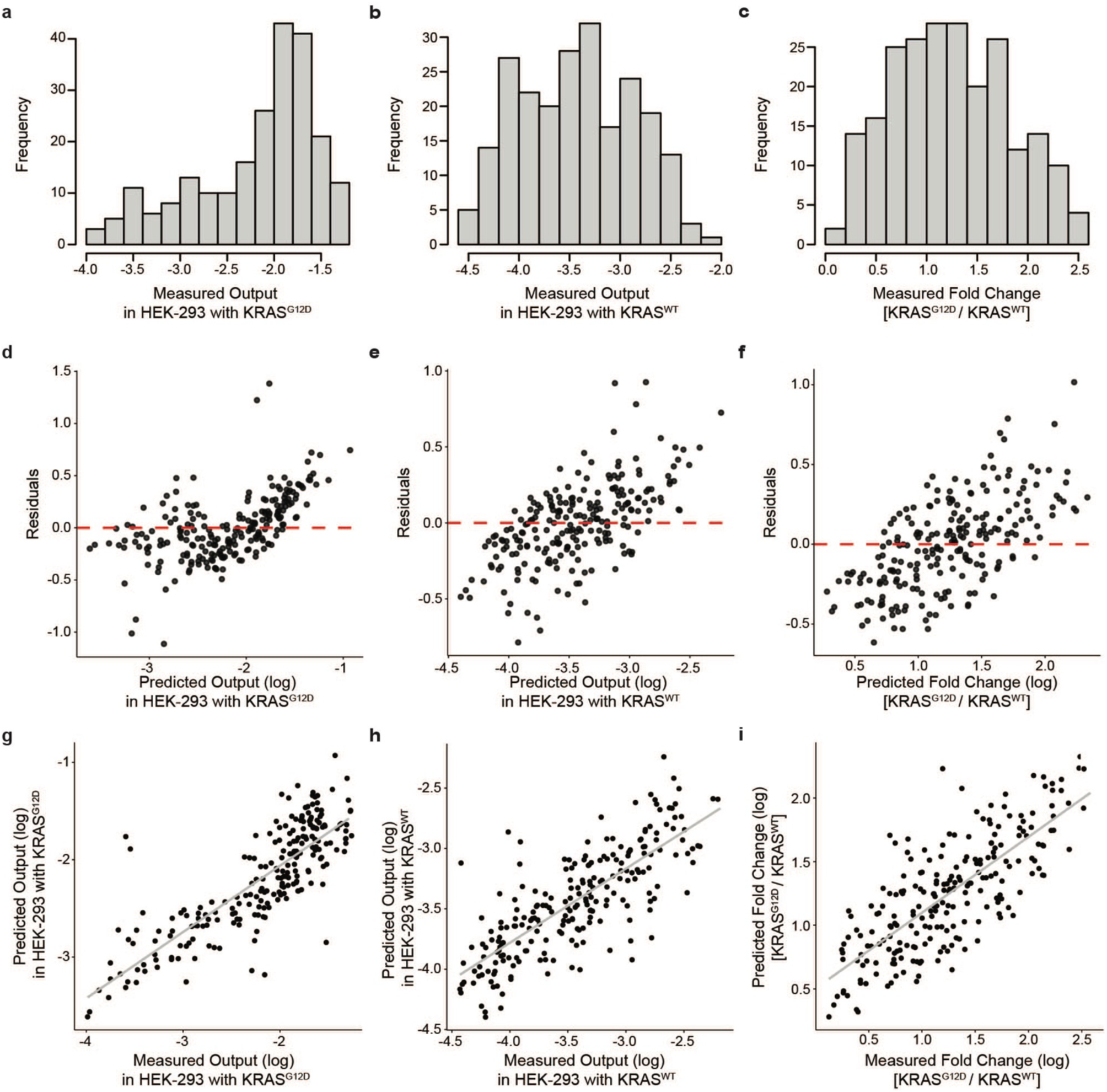
Assessment of the regression model (log-transformed model used in. Fig.5). (**a-c**) Histograms showing the distribution of the measured mCerulean output expression of all circuits tested in the screening in HEK293 overexpressing KRAS^G12D^ in **a**, KRAS^WT^ in **b**, or the fold change between KRAS^G12D^ and KRAS^WT^ in **c**. (**d-f**) Residuals plots showing the difference between measured and predicted output (residuals) vs the predicted output of HEK293 overexpressing KRAS^G12D^ in **d**, KRAS^WT^ in **e**, or the fold change between KRAS^G12D^ and KRAS^WT^ in **f** used to assess linearity and homoscedasticity assumptions. (**g-i**) Goodness of fit: Measurements of output vs predictions of the regression models based on the parts used in the circuit. Predicted values were calculated using the following general linear regression models. **a,d,g** show the ON-state data in HEK293 co-transfected with KRAS^G12D^. Model formula: log(RASmut_RU) ∼ RASmut_mCherry_AU + BD + RE_NarL_TAD + Linker + RE_NarX + Conc_NarX + Conc_NarL. **b,e,h** show the OFF-state in HEK293 co-transfected with KRAS^WT^. Model formula: log(RASwt_RU) ∼ RASwt_mCherry_AU + BD + RE_NarL_TAD + Linker + RE_NarX + Conc_NarX + Conc_NarL. **c,f,i** show the dynamic range calculated as fold change between ON- and OFF-state. Model formula: log(Ratio) ∼ RASmut_mCherry_AU + RASwt_mCherry_AU + BD + Linker + RE_NarX + RE_NarL + Conc_NarX + Conc_NarL Gray lines in g-I display the linear best fit used for calculating the Pearson correlation. RASmut_RU: mCerulean signal with KRAS^G12D^ normalized to transfection control, RASwt_RU: mCerulean signal with KRAS^WT^ normalized to transfection control, Ratio: fold change between RASmut_RU & RAS_wt_RU, RASmut_mCherry_AU: absolute transfection signal with KRAS^G12D^, RASmut_mCherry_AU: absolute transfection signal with KRAS^WT^, BD: binding domain, Linker: linker used in NarX fusion protein, RE_NarL_TAD: response element used to express NarL and transactivation domain fused to NarL, RE_NarX: response element used to express NarX proteins. Conc_NarX: transfected amount of plasmid expressing the NarX proteins. Conc_NarL: transfected amount of plasmid expressing the of NarL protein

**Supplementary Figure 17.**
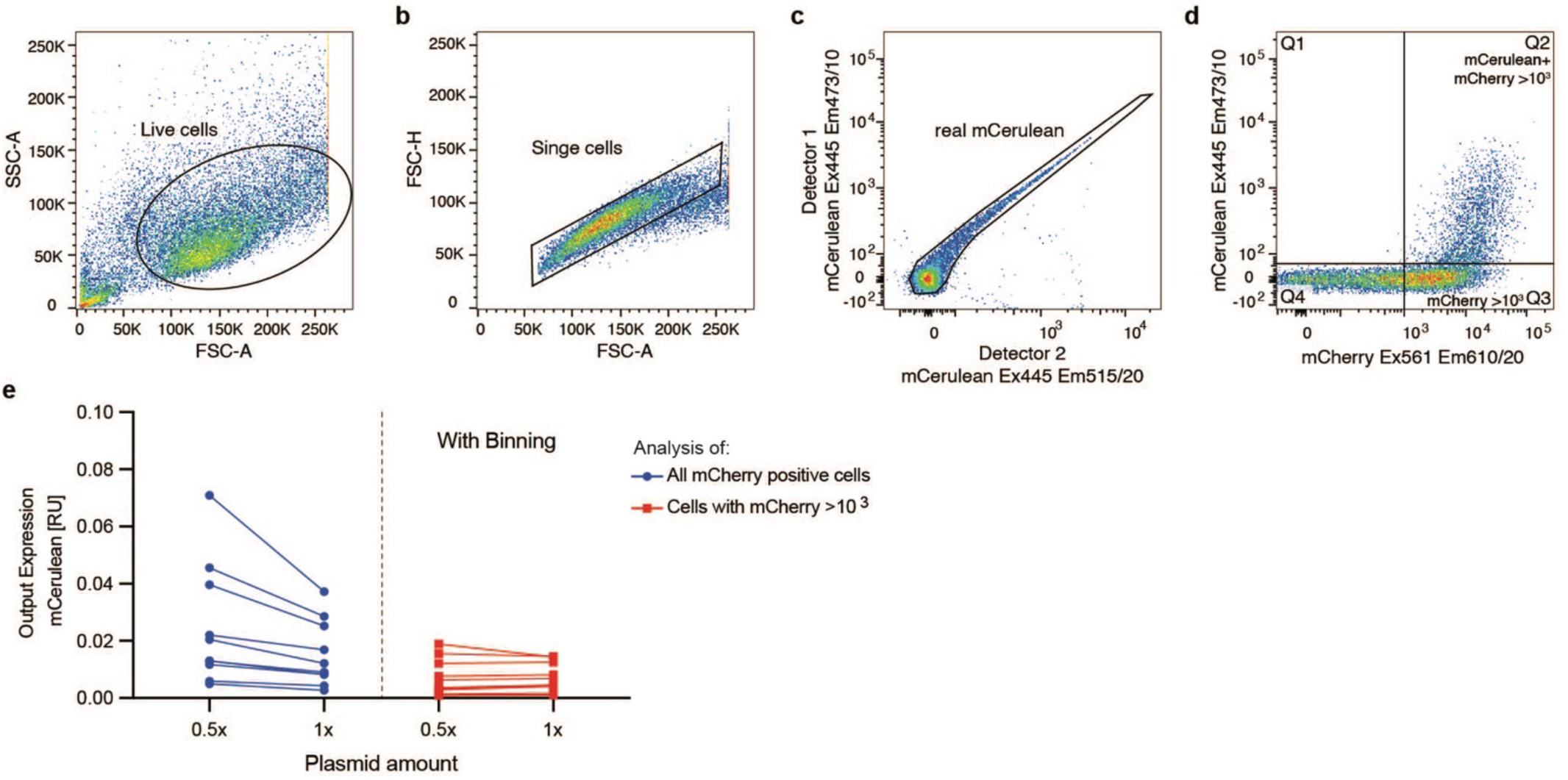
Illustration of gating in flow cytometry analyses. Gating example in HEK293 cells transfected with a mCerulean expressing RAS circuit and mCherry transfection control (**a**) Live cells are gated by plotting all events on a side scatter area (SSC-A) versus forward scatter area (FSC-A) density plot. (**b**) Single cells are gated by plotting FSC-A versus FSC-height (FSC-H). (**c**) Real mCerulean signals are separated from false positive signals by plotting the mCerulean signal measured with two independent detectors against each other. Both measure the signal from the 445 nm excitation laser, only exiting mCerulean and not mCherry. Both have an emission filter that can strongly detect mCerulean. Detector one measures mCerulean with a 473/11 emission filter, while detector two measures mCerulean with a 515/20 emission filter. Only real mCerulean signals where the signal correlates between the detectors are inside the gate. (**d**) Plot showing mCerulean output signal (excitation at 445 nm, emission filter 473/10) versus mCherry transfection control signal (excitation at 561 nm, emission filter 610/20). mCerulean output expression is calculated by multiplying frequency of parent * mean of Q2, which represents mCerulean positive cells with high transfection (mCherry higher than 10^3^). The mCherry signal is calculated by multiplying frequency of parent * mean of all cells with mCherry higher than 10^3^ (Q2+Q3). 10^3^ was chosen as threshold for transfection efficiency as cells below this rarely show circuit activation. (**e**) Binning makes the normalized Output signal less sensitive to different transfection amounts. Change in mCerulean output expression in HCT-116 cells transfected with 1x or 0.5x the plasmid amount of various RAS circuits to simulate different transfection efficiencies. All mCherry-positive cells were analyzed on the left (blue), while on the right the cells were binned for high transfection efficiency, and only cells with mCherry signal >10^3^ were analyzed (red).

**Supplementary Figure 18.**
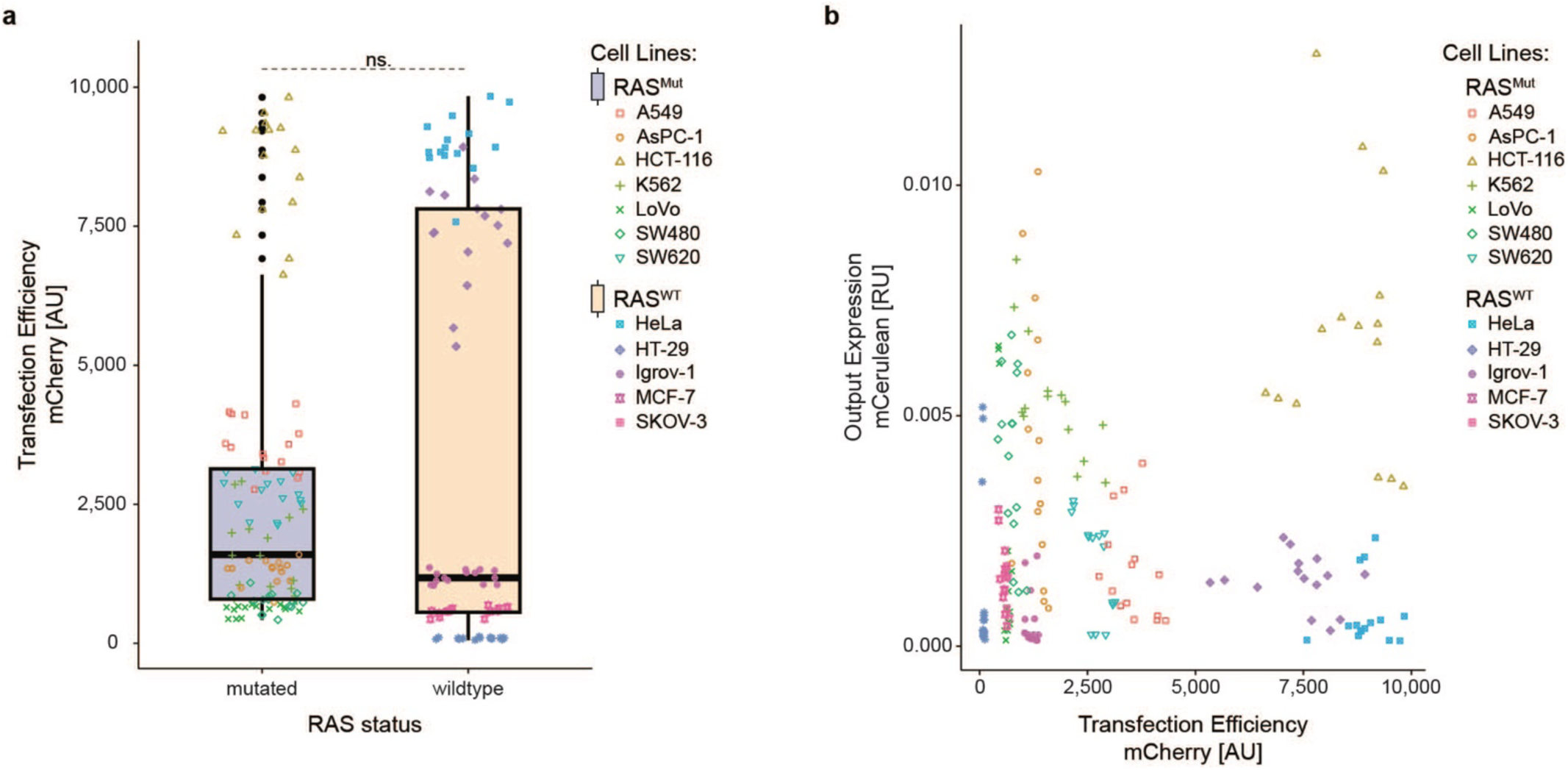
Correlation of Output Expression and Transfection Efficiency. (**a**) Comparison of transfection efficiency in cancer cell lines with (blue) or without (orange) mutation overactivating RAS. (**b**) Correlation between normalized output expression and transfection efficiency in the different cancer cell lines. mCerulean output expression and mCherry transfection efficiency were measured using flow cytometry. Absolute fluorescence units (AU) were calculated as mean*frequency of parent. mCerulean output expression AU was normalized to the mCherry transfection control AU to get the relative fluorescence units [RU]. Both **a & b** contain all samples tested during the cell line screening. Significance was tested using an unpaired two-tailed Student’s t-test.

## Supplementary Tables

**Supplementary Table 3:** Cell line and culturing details.

| Cell Line | Medium | Spitting Ratio | Provider | Catalogue Number | RRID | RAS status<sup>1</sup> |
| --- | --- | --- | --- | --- | --- | --- |
| **A-549** | RPMI-1640 | 1:2 - 1:15 | DSMZ | ACC 107 | CVCL_0023 | KRAS<sup>G12S</sup> (homozygous) |
| **AsPC-1** | RPMI-1640 | 1:2 - 1:6 | ATCC | CRL-1682 | CVCL_0152 | KRAS<sup>G12D</sup> (homozygous) |
| **HCT-116** | McCoy`s 5A | 1:2 - 1:20 | Abcam | ab288559 | CVCL_0291 | KRAS<sup>G13D</sup> (heterozygous) |
| **HCT-116**<br>**KRAS**<br>**knock out** | McCoy`s 5A | 1:2 - 1:15 | Abcam | ab276083 | CVCL_B1DS | KRAS knock out |
| **HEK293** | DMEM | 1:2 - 1:20 | Invitrogen | 11631-017 | CVCL_0045 | wildtype |
| **HeLa** | DMEM | 1:2 - 1:6 | ATCC | CCL-2 | CVCL_0030 | wildtype |
| **HT-29** | McCoy`s 5A | 1:2 - 1:8 | ATCC | HTB-38 | CVCL_0320 | Wildtype RAS; BRAF<sup>V600E</sup> (heterozygous) |
| **Igrov-1** | DMEM | 1:2 - 1:6 | Sigma-Aldrich | SCC203 | CVCL_1304 | wildtype |
| **K-562** | RPMI-1640 | 10<sup>5</sup> - 10<sup>6</sup> cells per mL | ATCC | CCL-243 | CVCL_0004 | BCR-ABL |
| **LoVo** | F-12K | 1:2-1:10 | ATCC | CCL-229 | CVCL_0399 | KRAS<sup>G13A</sup> (heterozygous) |
| **MCF-7** | DMEM | 1:2-1:6 | ATCC | HTB-22 | CVCL_0031 | wildtype |
| **SKOV-3** | McCoy`s 5A | 1:2-1:6 | ATCC | HTB-77 | CVCL_0532 | wildtype |
| **SW480** | DMEM | 1:2 - 1:8 | ATCC | CCL-228 | CVCL_0546 | KRAS<sup>G12V</sup> (homozygous) |
| **SW620** | DMEM | 1:2 - 1:10 | ATCC | CCL-227 | CVCL_0547 | KRAS<sup>G12V</sup> (homozygous) |

<sup>1</sup>Bairoch A. The Cellosaurus, a cell line knowledge resource. J. Biomol. Tech. 29:25-38 (2018).
DOI: 10.7171/jbt.18-2902-002; PMID: 29805321; PMCID: PMC5945021

**Supplementary Table 6:**
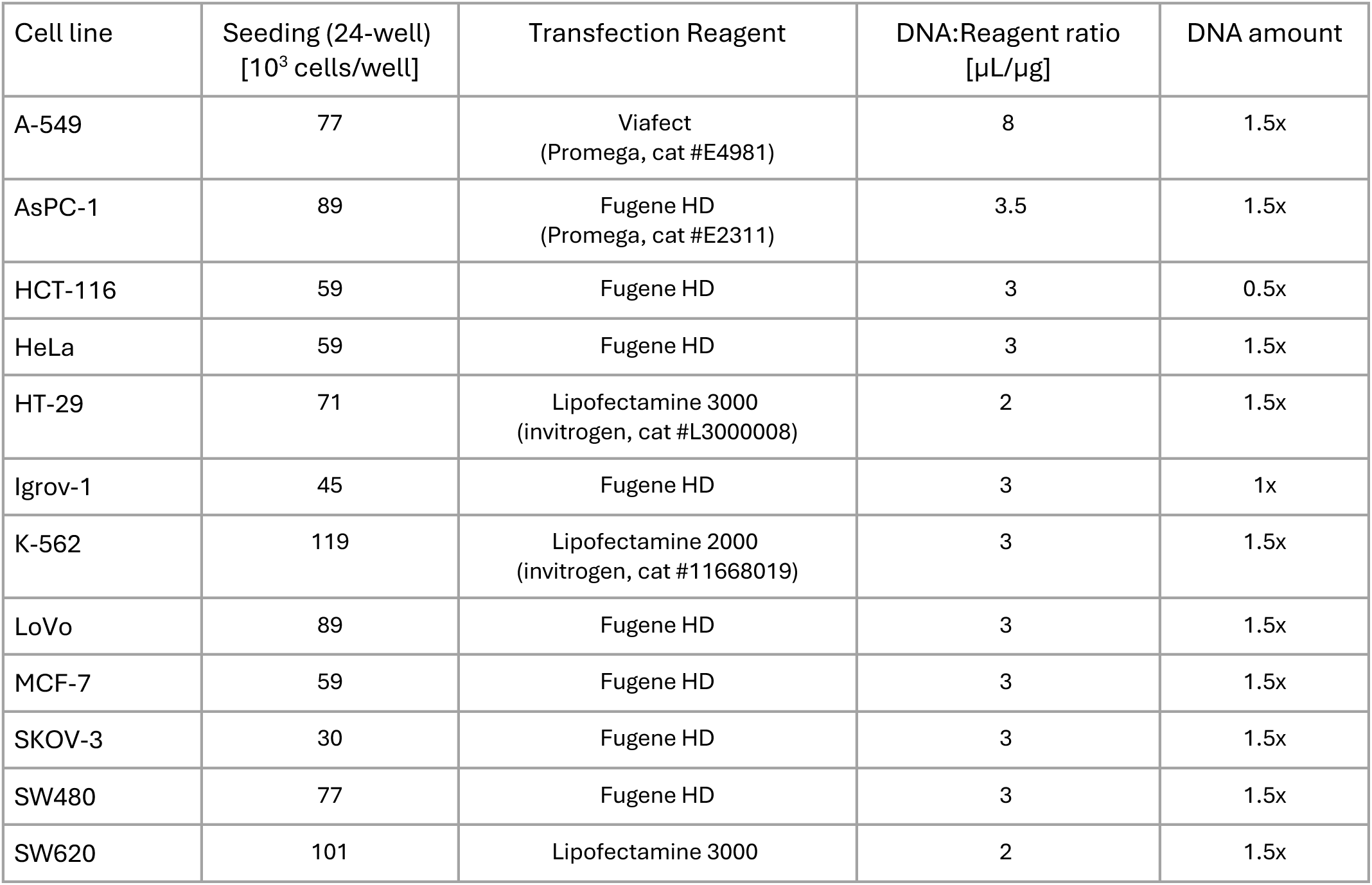
Seeding and transfection conditions in different cancer cell lines (Fig.7 & 8d-l)

| Cell line | Seeding (24-well)<br>[10<sup>3</sup> cells/well] | Transfection Reagent | DNA:Reagent ratio<br>[μL/μg] | DNA amount |
| --- | --- | --- | --- | --- |
| A-549 | 77 | Viafect<br>(Promega, cat #E4981) | 8 | 1.5x |
| AsPC-1 | 89 | Fugene HD<br>(Promega, cat #E2311) | 3.5 | 1.5x |
| HCT-116 | 59 | Fugene HD | 3 | 0.5x |
| HeLa | 59 | Fugene HD | 3 | 1.5x |
| HT-29 | 71 | Lipofectamine 3000<br>(invitrogen, cat #L3000008) | 2 | 1.5x |
| Igrov-1 | 45 | Fugene HD | 3 | 1x |
| K-562 | 119 | Lipofectamine 2000<br>(invitrogen, cat #11668019) | 3 | 1.5x |
| LoVo | 89 | Fugene HD | 3 | 1.5x |
| MCF-7 | 59 | Fugene HD | 3 | 1.5x |
| SKOV-3 | 30 | Fugene HD | 3 | 1.5x |
| SW480 | 77 | Fugene HD | 3 | 1.5x |
| SW620 | 101 | Lipofectamine 3000 | 2 | 1.5x |

**Supplementary Table 7:** Seeding and transfection conditions for all experiments (Fig.1-6e) except the cancer cell line screening (Fig.7)

| Cell line | Seeding<br>96-well<br>[10<sup>3</sup> cells/<br>well] | Seeding<br>24-well<br>[10<sup>3</sup> cells/<br>well] | Seeding<br>6-well<br>[10<sup>3</sup> cells/<br>well] | Transfection<br>Reagent | DNA:Reagent ratio<br>[μg/μL] | DNA amount |
| --- | --- | --- | --- | --- | --- | --- |
| HEK293 | 20 | 75 | 400 | Lipofectamine 2000 | 2 | 1x |
| HCT-116<br>wildtype | 10 | 60 | - | Fugene | 3 | 0.25-1x* |
| HCT-116<br>KRAS k.o. | 20 | 100 | - | Fugene | 3 | 1x |

\*see also Supplementary Table S4

**Supplementary Table 8:** Excitation filter, dichroic mirror, and emission filter wavelengths used during fluorescence microscopy.

| Fluorochrome | Excitation Filter [nm] | Dichroic Mirror [nm] | Emission Filter [nm] | Exposure Times [ms] |
| --- | --- | --- | --- | --- |
| mCerulean | 438/24 | 458 | 483/32 | 500 |
| mCherry / mScarlet | 562/40 | 593 | 624/40 | 300 |
| SBFP2 | 370/36 | 495 | 520/35 | 300 |

**Supplementary Table 9:** Excitation laser, long pass filter, and emission filter wavelengths used during flow cytometry.

| Fluorochrome | Excitation Laser [nm] | Long pass Filter [nm] | Emission Filter [nm] |
| --- | --- | --- | --- |
| mCerulean | 445 | - | 473/10 |
| SYFP2 / mVenus | 488 | 505 | 530/11 |
| mCherry / mScarlet | 561 | 600 | 610/20 |
| SBFP2 | 405 | - | 445/15 |

**Supplementary Table 10:** Tested regression models to select the independent variables and assess the interaction between response element and expressed parts.

|  | Model 1 | Model 2 | Model 3 (Selected Model) |
| --- | --- | --- | --- |
| Independent Variable(s) used to describe the response element (RE) | RE | RE_NarX + RE_NarL | RE_NarX + RE_NarL_TAD |
| Description/ Assumptions | The effect of the response elements is independent of what part is expressed. | The effect of response elements depends on whether they are used to express the NarX fusion proteins (RE_NarX) or NarL protein (RE_NarL). | The effect of response elements depends on whether they are used to express the NarX fusion proteins or NarL protein. Additionally, the effect of REs depends on the TAD fused to NarL (RE_NarL_TAD). |
| Complete Formula used in regression model (ON-State Model) | $\text{Log(RASmut\_RU)} \sim \text{RASmut\_mCherry} + \text{RE} + \text{BD} + \text{TAD} + \text{Linker} + \text{Conc\_NarX} + \text{Conc\_NarL}$ | $\text{Log(RASmut\_RU)} \sim \text{RASmut\_mCherry} + \text{RE\_NarX} + \text{RE\_NarL} + \text{BD} + \text{TAD} + \text{Linker} + \text{Conc\_NarX} + \text{Conc\_NarL}$ | $\text{Log(RASmut\_RU)} \sim \text{RASmut\_mCherry} + \text{RE\_NarX} + \text{RE\_NarL\_TAD} + \text{BD} + \text{Linker} + \text{Conc\_NarX} + \text{Conc\_NarL}$ |
| ON-State Model (HEK<sup>G12D</sup>): | Pearson R<sup>2</sup> = 0.44<br>Spearman = 0.62 | Pearson R<sup>2</sup> = 0.45<br>Spearman = 0.63 | Pearson R<sup>2</sup> = 0.68<br>Spearman = 0.82 |
| OFF-State Model (HEK<sup>WT</sup>): | Pearson R<sup>2</sup> = 0.44<br>Spearman = 0.67 | Pearson R<sup>2</sup> = 0.53<br>Spearman = 0.73 | Pearson R<sup>2</sup> = 0.61<br>Spearman = 0.78 |
| Dynamic Range Model (HEK<sup>G12D</sup>/HEK<sup>WT</sup>): | Pearson R<sup>2</sup> = 0.44<br>Spearman = 0.66 | Pearson R<sup>2</sup> = 0.50<br>Spearman = 0.70 | Pearson R<sup>2</sup> = 0.60<br>Spearman = 0.75 |

**Supplementary Table 11:** Numerical values from flow cytometry histograms in Fig.5d.

| Circuit ID | Results in HEK293 with KRAS<sup>G12D</sup><br>(Results from concatenated triplicates) |  |  | Results in HEK293 with KRAS<sup>WT</sup><br>(Results from concatenated triplicates) |  |  |
| --- | --- | --- | --- | --- | --- | --- |
|  | Mean mCerulean expression | Freq. of Parent [% of all measured cells] | Number of mCerulean positive cells | Mean mCerulean expression | Freq. of Parent [% of all measured cells] | Number of mCerulean positive cells |
| RAS_Sensor_F.L.T. | 1361 | 1.9 | 666 | 1013 | 0.5 | 119 |
| PY2_NarL-F.L.T. | 1463 | 4.2 | 1456 | 93 | 0.2 | 14 |
| PY2_NarX_F.L.T. | 1763 | 4.0 | 1587 | 297 | 0.2 | 24 |
| PY2_all_F.L.T. | 1707 | 7.1 | 2547 | 191 | 0.3 | 26 |

**Supplementary Table 12:**
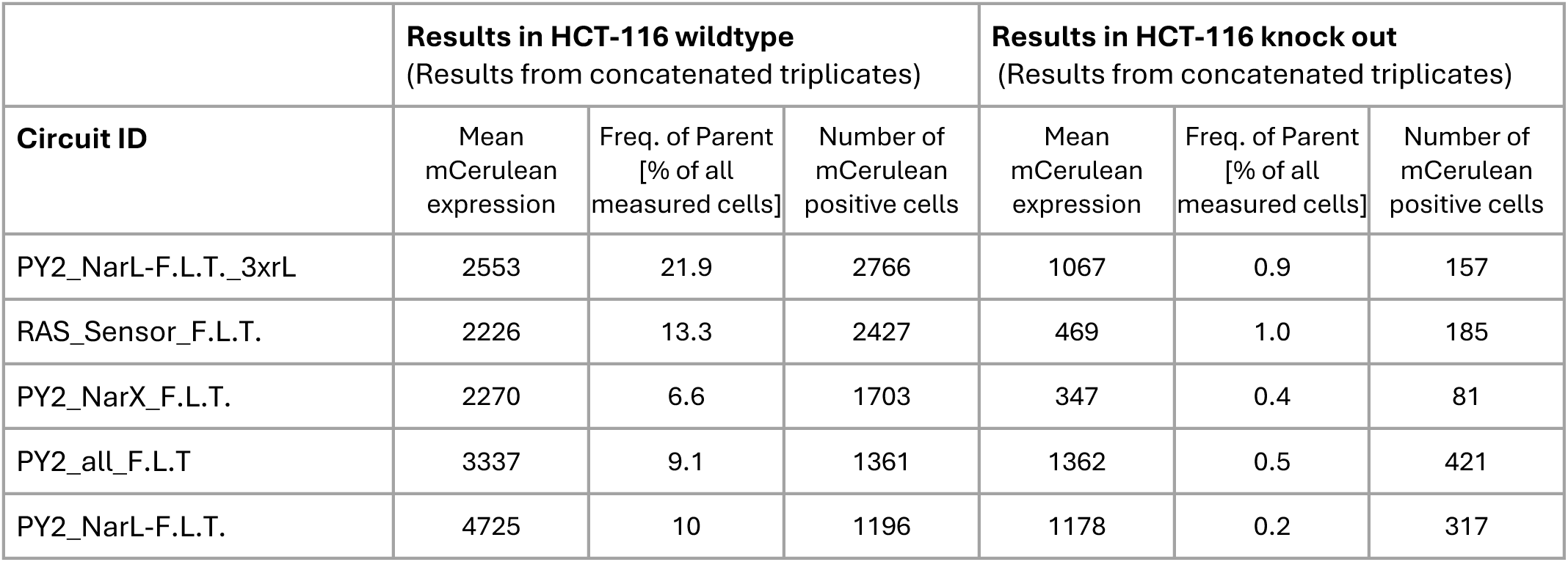
Numerical values from flow cytometry histograms in Fig.6e.

| **Circuit ID** | **Results in HCT-116 wildtype**<br>(Results from concatenated triplicates) |  |  | **Results in HCT-116 knock out**<br>(Results from concatenated triplicates) |  |  |
| --- | --- | --- | --- | --- | --- | --- |
|  | Mean mCerulean expression | Freq. of Parent [% of all measured cells] | Number of mCerulean positive cells | Mean mCerulean expression | Freq. of Parent [% of all measured cells] | Number of mCerulean positive cells |
| PY2_NarL-F.L.T._3xrL | 2553 | 21.9 | 2766 | 1067 | 0.9 | 157 |
| RAS_Sensor_F.L.T. | 2226 | 13.3 | 2427 | 469 | 1.0 | 185 |
| PY2_NarX_F.L.T. | 2270 | 6.6 | 1703 | 347 | 0.4 | 81 |
| PY2_all_F.L.T | 3337 | 9.1 | 1361 | 1362 | 0.5 | 421 |
| PY2_NarL-F.L.T. | 4725 | 10 | 1196 | 1178 | 0.2 | 317 |

**Supplementary Table 13:** Segmentation parameters for image analysis using Agilent eSight in Fig.8.

| Cell Line – Channel | Segmentation Adjustment<br>(Background <-> Cell) | Cleanup (Hole Fill) | Area Filter<br>(Minimal Cell Size) | Edge Sensitivity |
| --- | --- | --- | --- | --- |
| HCT-116 – Bright field | 0.6 | 0 | 1000 | - |
| HCT-116 – Red Fluorescence | 0.6 | 500 | 100 | 0 |
| Igrov-1 – Bright field | 0.9 | 0 | 1500 | - |
| Igrov-1 – Red Fluorescence | 1.3 | 500 | 100 | 0 |
| SW620 – Bright field | 0.9 | 0 | 1000 | - |
| SW620 – Red Fluorescence | 1 | 500 | 100 | 0 |

## Notes

### Summary of Updates

1) We validated the generalizability of the RAS sensor by comparing its activation upon transfection with various RAS mutants and isoforms in HEK293 cells. We confirmed robust activation across oncogenic variants, presented in Figure 1g and Supplementary Fig. 2. 2) To assess the functionality of the RAS-targeting circuits to kill cancer cells, we replaced the mCerulean on the output plasmid with a clinically relevant suicide gene, herpes simplex virus thymidine kinase (HSV-TK), which converts ganciclovir into a cytotoxic compound. We performed time-course killing assays in two RAS-mutant lines (HCT-116 and SW620) and one wild-type line (Igrov-1). At high circuit doses, we observed robust killing of transfected and bystander HCT-116 cells. At lower doses (HCT-116) or in cells with lower transfection efficiency (SW620), only transfected cells were killed. In contrast, Igrov-1 cells continued to grow, indicating preferential cytotoxicity in RAS-mutant backgrounds. While differences between the cell lines may influence the circuit-mediated killing and limit a definitive conclusion regarding selectivity, the data demonstrates that RAS-targeting circuits can be used to efficiently kill RAS-driven cancer cell lines and supports a preferential cytotoxic effect in RAS-mutant cell lines. These results are now presented as a new chapter (Fig. 8 and Supplementary Fig. 14). 3) Additionally, we further investigated the mechanism behind the observed RAS-dependent output expression, focusing on potential contributions from EF1α-driven sensor overexpression and non-linear amplification via the NarX-NarL two-component system (TCS). We cloned and tested both membrane-bound and cytosolic constitutively active NarX variants alongside various negative controls. While we observed non-linear amplification of the output, we found that this is not the full explanation for the RAS-dependent increase in output expression and confirmed that RAS-binding is required for full activation of the RAS sensor by RAS (Supplementary Fig.4). To assess the contribution of response elements, we compared RAS-targeting circuits (Fig. 4f) to controls expressing constitutively active NarX without RAS binding. Expressing constitutively active NarX instead of the RAS-binding fusion proteins showed reduced dynamic ranges between KRASG12D and KRASWT cells in all but one condition, highlighting that combining MAPK response elements with the RAS-binding sensor increases maximal fold changes (Supplementary Fig. 7).

